# Terminal Schwann Cells Are Required for Presynaptic Vesicle Homeostasis but Not for Gross Neuromuscular Junction Architecture in Mice

**DOI:** 10.64898/2026.06.15.732329

**Authors:** Hyun Kim, Seon-Yong Kim, Kyusang Yoo, Se-Young Choi, Young-Yun Kong

**Affiliations:** School of Biological Sciences, Seoul National University, Seoul 08826, Republic of Korea; Department of Physiology, Dental Research Institute, Seoul National University School of Dentistry, Seoul 03080, Republic of Korea

**Keywords:** Neuromuscular junction, terminal Schwann cell, *Col20a1*, synaptic transmission, vesicle homeostasis

## Abstract

Terminal Schwann cells (tSCs) are specialized glial cells persistently residing at the neuromuscular junction (NMJ), where they are widely held to participate in synaptic transmission and maintenance. However, elucidating their precise *in vivo* roles has been hindered by a lack of models for tSC-specific genetic manipulation, as conventional tools simultaneously targeted axonal Schwann cells. Here, we identify *Col20a1* as a gene highly specific to tSCs through single-cell transcriptomic analysis and generate a novel *Col20a1*-CreERT2 knock-in mouse model. Utilizing this model, we achieved systemic labeling and ablation of tSCs during the postnatal maturation window. Surprisingly, systemic tSC loss disrupted neither gross NMJ architecture nor overall motor behavior. Instead, electrophysiological and ultrastructural analyses revealed presynaptic defects, characterized by increased frequency and reduced amplitude of spontaneous miniature endplate potentials (mEPPs), accelerated synaptic depression during high-frequency stimulation, and reduction of presynaptic vesicle density. Together, our findings establish *Col20a1* as a genetic handle for tSC research and show that tSCs are required for presynaptic vesicle homeostasis rather than maintaining the gross architecture of the synapse.

## Introduction

The neuromuscular junction (NMJ) is a highly specialized peripheral synapse that transmits motor signals from neurons to skeletal muscles. Because of this pivotal role, impairment in NMJ integrity or function is linked to various neuromuscular disorders, such as myasthenia gravis (Darabid et al., 2018; Engel et al., 2003; Plomp et al., 2015), amyotrophic lateral sclerosis (ALS) (De Winter et al., 2006; Dupuis & Loeffler, 2009; Khabibrakhmanov et al., 2025), and spinal muscular atrophy (SMA) (Kariya et al., 2008; Kong et al., 2009). Thus, investigating the precise mechanisms underlying NMJ function is of critical importance. In addition to the presynaptic motor nerve terminal and the postsynaptic muscle fiber, the NMJ has long been recognized as a tripartite synapse that incorporates a dedicated glial compartment, the terminal Schwann cells (tSCs) (Araque et al., 2014; Araque et al., 1999; Barik et al., 2016; Kettenmann & Ransom, 2012; Ko & Robitaille, 2015; Sanes & Lichtman, 2001; Santosa et al., 2018; Todd et al., 2007; Volterra et al., 2002; Wu et al., 2010). Decades of work have established that tSCs monitor and modulate synaptic activity and contribute to the formation, maintenance, and repair of the synapse (Barik et al., 2016; Darabid et al., 2018; Fuertes-Alvarez & Izeta, 2021; Hastings & Valdez, 2024; Petrov et al., 2021). What has remained challenging is the ability to manipulate them selectively *in vivo*. Because tSCs share most canonical markers with the other Schwann cell subtypes of the peripheral nerve, experimental access to tSCs without concurrently affecting the axonal Schwann cells (aSCs) that ensheathe the motor axon has been difficult to achieve (Hastings & Valdez, 2024; Jablonka□Shariff et al., 2021; Santosa et al., 2018).

In mice, while the nascent NMJ structure forms before birth, the extensive NMJ maturation occurs within the first three postnatal weeks (Kariya et al., 2014; Medina Moreno & Henríquez, 2022; Sanes & Lichtman, 1999, 2001). NMJs transition from polyneuronal to mononeuronal innervation through competitive synapse elimination (Balice-Gordon & Lichtman, 1993; Brown et al., 1976; Sanes & Lichtman, 1999), and postsynaptic acetylcholine receptor (AChR) clusters mature from plaque-like structure into complex pretzel-like structure, accompanied by the extensive formation of junctional folds (Marques et al., 2000; Matthews-Bellinger & Salpeter, 1983; Shi et al., 2012). During this developmental window, tSCs support nerve pruning process by phagocytosing axosomes (Bishop et al., 2004; Darabid et al., 2013; Lee et al., 2016; Riley, 1981; Smith et al., 2013) and decode synaptic activity through Ca^2+^ and purinergic signaling (Darabid et al., 2013; Darabid et al., 2018; Heredia et al., 2018; Jahromi et al., 1992; Reist & Smith, 1992; Robitaille, 1998). The fundamental necessity of Schwann cells has been further evidenced by studies using ErbB2 or ErbB3 mutant mice, where the failure of Schwann cell formation and migration led to severe nerve retraction (Lin et al., 2000; Morris et al., 1999; Riethmacher et al., 1997; Woldeyesus et al., 1999). Furthermore, ablating tSCs in frog NMJ by antibody and complement system reduced nerve growth and induced nerve retraction, suggesting that tSCs are vital for long-term maintenance of NMJ (Feng & Ko, 2008; Koirala et al., 2003). Whether the mammalian NMJ depends on tSCs to a comparable degree, however, has been difficult to test directly.

Efforts to define tSC function in mammals have long relied on drivers expressed throughout the Schwann cell lineage, such as *Plp1* and *S100b* (Barik et al., 2016; Zuo et al., 2004). Because these drivers also act on the aSCs that ensheathe the motor axon, phenotypes obtained with them require careful interpretation to separate tSC-specific contributions from those of aSCs (Barik et al., 2016). Marker combinations have since narrowed this gap: *S100b*-GFP;NG2-dsRed double-positive cells label tSCs selectively within skeletal muscle, allowing these cells to be visualized and isolated (Castro et al., 2020). Such dual-reporter strategies, however, address identification rather than manipulation, and do not by themselves provide a route to Cre-dependent labeling or ablation. *Col20a1* was identified among the genes enriched in this population (Castro et al., 2020), and its loss in Schwann cells alters NMJ regeneration and locomotor activity (Hastings et al., 2025). The locus itself, however, has not been used to gain inducible genetic access to tSCs. What has been missing is a single-locus, inducible driver for genetic manipulation in tSCs, which would render the cell type itself experimentally accessible.

Here, we establish Col20a1 as a genetically tractable marker for tSCs. We first confirmed and extended its specificity by applying two independent selection criteria to skeletal muscle single-cell transcriptomes and by cross-validating the result against independent bulk RNA-seq datasets, which together distinguished Col20a1 from markers of myelinating Schwann cells (mSCs) and Remak non-myelinating Schwann cells (nmSCs). We then generated a Col20a1-CreERT2 knock-in mouse line, which achieved labeling and diphtheria toxin-mediated ablation of tSCs throughout the body while sparing aSCs. We induced ablation at postnatal day 10 (P10), during the later phase of postnatal NMJ maturation, when the synapse undergoes structural refinement and acquires its adult functional properties. Within this window, systemic tSC ablation did not produce overt disruption of gross NMJ architecture or of motor behavior. Electrophysiological and ultrastructural analyses, however, revealed a selective requirement for tSCs in sustaining presynaptic vesicle homeostasis and in maintaining synaptic efficacy during sustained high-frequency transmission. Our study thus provides a genetic platform for studying tSC function *in vivo* and refines the contribution of these cells to the mammalian NMJ.

## Results

### Selection of *Col20a1* as a specific marker for tSCs

To develop a tSC-specific genetic model, we needed to identify a reliable tSC-specific marker. To this end, we analyzed gene expression profiles of muscle resident cells to find genes highly and specifically expressed in tSCs. To investigate the transcriptomic profile of Schwann cell populations residing in skeletal muscle, we analyzed a publicly available single-cell RNA sequencing (scRNA-seq) dataset of skeletal muscle-resident cells (Giordani et al., 2019). We first generated a UMAP plot of the entire muscle cell population (Figure 1A) and identified the Schwann cell cluster based on the expression of canonical glial markers *Plp1* and *S100b* (Figure 1B; Figure 1—figure supplement 1). Subsequent isolation and sub-clustering of this Schwann cell population revealed three distinct clusters (Figure 1C). To annotate the identity of each sub-cluster, we evaluated the expression of well-known marker genes for Schwann cell subtypes. Cluster 1 and Cluster 2 exhibited high expression of myelinating Schwann cell (mSC) and Remak non-myelinating Schwann cell (nmSC) markers, respectively. Notably, Cluster 3 displayed significant enrichment for genes previously described to be expressed in tSCs (Castro et al., 2020), indicating that Cluster 3 represents the tSC population (Figure 1D; Figure 1—figure supplement 2). Clusters 1 and 2 likely correspond to the glial populations residing within intramuscular nerve bundles, representing mSCs and nmSCs that envelop the axons. The mSC population (Cluster 1) is also presumed to include myelinating axonal Schwann cells (aSCs) that wrap the motor axons leading to the NMJ. The anatomical distribution of these defined Schwann cell subtypes within skeletal muscle is schematized in Figure 1E.

**Figure 1.**
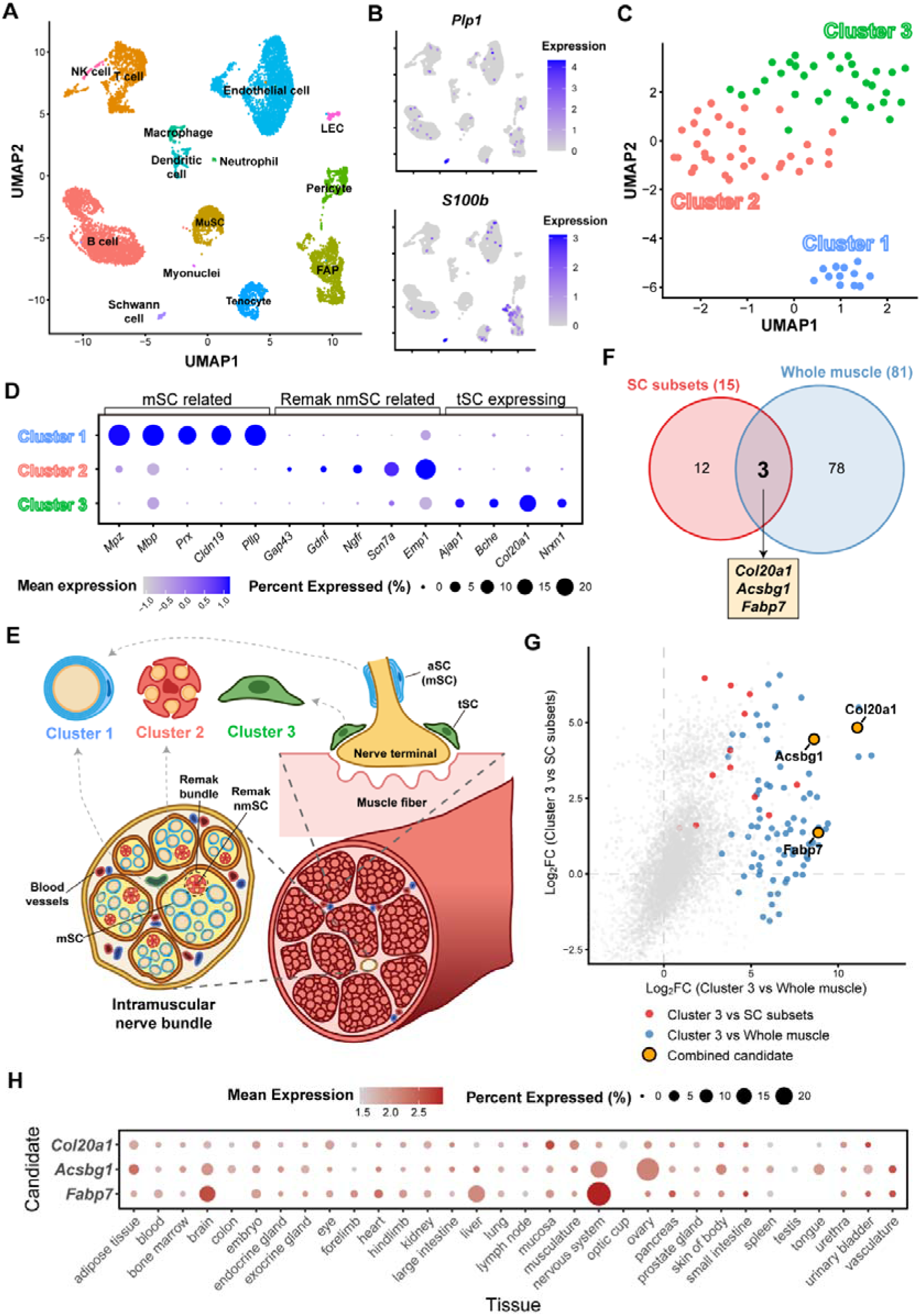
Selection of *Col20a1* as a specific marker for terminal Schwann cells (tSCs) in skeletal muscle. (A) UMAP plot of the entire skeletal muscle-resident cell populations from a public scRNA-seq dataset (Giordani et al., 2019). (B) Feature plots showing the expression of canonical Schwann cell markers, *Plp1* and *S100b*. (C) Sub-clustering of the isolated Schwann cell population reveals three distinct sub-populations (Clusters 1–3). (D) Dot plot displaying the expression profiles of established marker genes for myelinating Schwann cells (mSC, Cluster 1), Remak non-myelinating Schwann cells (Remak nmSC, Cluster 2), and tSC-expressing genes (Cluster 3). Node size indicates the percentage of cells expressing the gene, and color intensity represents the mean expression level. (E) Schematic illustration depicting the anatomical distribution of the three defined Schwann cell subtypes within the skeletal muscle and neuromuscular junction. (F) Venn diagram illustrating the intersection of two differential gene expression screening criteria: genes highly expressed in Cluster 3 compared to other SC subsets (15 genes; log_2_FC > 1, adjusted *p*-value < 0.05, pct.1 > 0.5), and genes specifically expressed in Cluster 3 compared to all other muscle cells (81 candidates; log_2_FC > 1, adjusted *p*-value < 0.05, pct.2 < 0.01). Three shared candidate genes (*Col20a1*, *Acsbg1*, and *Fabp7*) were identified. (G) Scatter plot comparing the Log2 fold-change (FC) of gene expression for Cluster 3 vs SC subsets (y-axis) and Cluster 3 vs whole muscle cells (x-axis). Genes that passed each screening criterion are depicted by red and blue dots, respectively. The intersected candidate genes are highlighted in an orange circle. (H) Dot plot from the CELLxGENE database (Program et al., 2025) showing the cross-tissue expression profiles of the three candidate genes, highlighting the minimal expression of *Col20a1* in the central nervous system compared to *Acsbg1* and *Fabp7*.

Next, we sought to identify a highly specific marker gene for tSCs by performing rigorous differentially expressed gene (DEG) analysis using two distinct criteria. First, to distinguish tSCs from other Schwann cell subtypes, we compared Cluster 3 against Clusters 1 and 2. We focused on candidates showing widespread expression within tSCs, selecting only those expressed in at least 50% of Cluster 3 cells with at least a twofold increase in expression relative to other Schwann cell populations, yielding 15 candidates. Second, to ensure specificity within the entire skeletal muscle tissue, we compared Cluster 3 against all other non-Schwann muscle cells. To guarantee that the markers are highly restricted to tSCs, we filtered for genes expressed in less than 1% of other cell populations, while also exhibiting at least a twofold increase in expression, which resulted in 81 candidates. The intersection of these two criteria identified only three candidate genes: *Col20a1*, *Acsbg1*, and *Fabp7* (Figure 1F, G; Figure 1—figure supplement 3).

To minimize potential off-target effects in systemic applications or genetic targeting models, we further scrutinized systemic expression profiles of these three candidate genes using the CELLxGENE database (Figure 1H) (Program et al., 2025). We noted that *Acsbg1* and *Fabp7* showed considerable expression in the brain and central nervous system. Since our objective is to investigate tSC function at the NMJ, we prioritized a marker with minimal expression in adjacent neuromuscular components to avoid artefacts in subsequent genetic manipulation models. *Col20a1* showed relatively minimal expression in these regions. While slight expression was observed in mucosal tissues, its overall restricted expression pattern makes *Col20a1* a favorable candidate for investigating tSC functions. Furthermore, re-analysis of independent bulk RNA-seq datasets (Castro et al., 2020; Ham et al., 2020) confirmed that *Col20a1* has the highest expression level and most significant fold-change among the three candidates in S100b^+^/NG2^+^ double-positive cells and NMJ-enriched regions (Figure 1—figure supplement 4). Taken together, *Col20a1* was highly prioritized in various criteria for an ideal tSC-specific marker in skeletal muscle, leading us to select it for the generation of tSC-specific genetic model.

Finally, we examined how *Col20a1* relates to markers already associated with tSC. *Cspg4* (NG2), which labels a subset of non-myelinating Schwann cells when combined with S100b (Castro et al., 2020), was expressed across the Schwann cell populations of skeletal muscle and, within them, was enriched in Cluster 3 (Figure 1—figure supplement 5A, B). Its expression therefore resolves to the same subpopulation as *Col20a1*. *Kcnj10* (Kir4.1), which has been reported in tSCs (Procacci et al., 2023), was not captured in the single-cell data, but was likewise enriched in the S100b^+^/NG2^+^ population and in NMJ-enriched regions of the bulk datasets (Figure 1—figure supplement 5C). These observations indicate that *Col20a1* marks a tSC population overlapping with that labeled by *Cspg4* and *Kcnj10*, rather than a separate subset, and that *Col20a1* can therefore serve as a marker for this population.

### Generation of *Col20a1-*CreERT2 mouse model and validation of its universal expression in tSCs

Building upon our identification of *Col20a1* as a highly specific tSC marker, we generated a *Col20a1-*CreERT2 knock-in mouse model to selectively target tSCs *in vivo*. P2A-CreERT2 cassette was inserted at the C-terminus of the endogenous *Col20a1* gene (Figure 2—figure supplement 1A). We crossed these mice with a *Rosa26*-LSL-tdTomato (*Rosa*-tdT) reporter line to generate *Col20a1-*CreERT2*; Rosa-*tdT (*Col20a1-*tdT) mice. To induce recombination, adult mice were administered tamoxifen via oral gavage (160 mg/kg body weight) for five consecutive days, and the extensor digitorum longus (EDL) muscles were analyzed one week after the first dose. Immunofluorescence analysis of the neuromuscular junctions revealed robust tdTomato expression specifically within the tSC region in *Col20a1-*tdT mice, whereas control *Rosa*-tdT mice lacking the CreERT2 allele showed no detectable signal (Figure 2A). High-magnification imaging further confirmed that the tdTomato signal co-localized tightly with the S100B glial marker and accurately enveloped the alpha-bungarotoxin (Btx)-labeled AChR region. DAPI staining confirmed the distinct cellular morphology of these labeled tSCs (Figure 2B).

**Figure 2.**
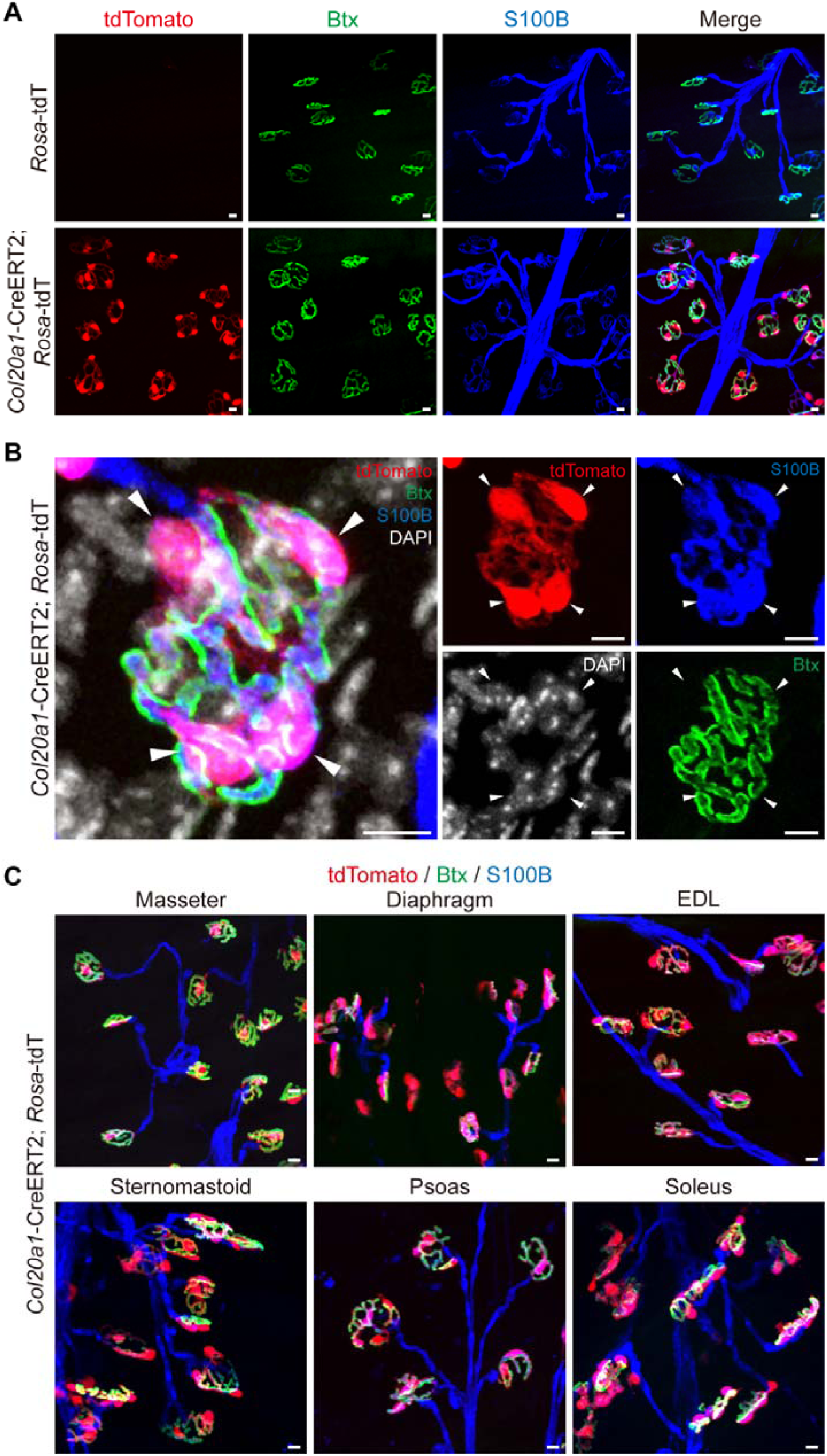
Generation of the *Col20a1*-CreERT2 mouse model and validation of its system-wide expression in tSCs. Adult mice (8−12 weeks old) were treated with tamoxifen (TMX) via oral gavage (160 mg/kg body weight) for five consecutive days and analyzed one week after the initial dose. (A) Representative immunofluorescence images of EDL muscle neuromuscular junctions (NMJs) from control (*Rosa*-tdT) and *Col20a1*-CreERT2; *Rosa*-tdT mice. The tdTomato reporter signal (red) is robustly detected at the NMJs of *Col20a1*-CreERT2; *Rosa*-tdT mice, and was not observed in control *Rosa*-tdT mice. Postsynaptic acetylcholine receptors (AChRs) are labeled with α-bungarotoxin (Btx, green), and glial cells are labeled with S100B (blue). (B) High-magnification confocal images highlighting the precise cellular localization of the tdTomato reporter. White arrowheads indicate the tight co-localization of tdTomato (red) with the tSC marker S100B (blue), which effectively envelops the postsynaptic AChR clusters (green). DAPI (white) staining indicates nuclei. (C) Representative immunofluorescence images showing the spatial distribution of the reporter across diverse anatomical locations, including cranial (masseter, sternomastoid), axial (diaphragm, psoas), and appendicular (EDL, soleus) muscles. The *Col20a1*-CreERT2; *Rosa*-tdT model reliably targets tSCs across all examined muscle types. Scale bars: 10 µm in all images.

To quantitatively validate the cell-type specificity of our model, we performed fluorescence-activated cell sorting (FACS) on whole muscle mononuclear cells. Cells were segregated into Lineage-positive (Lin^+^; CD31^+^, CD45^+^, or Ter119^+^), muscle stem cells (MuSCs; Lin^−^/VCAM1^+^/Sca-1^−^), fibro-adipogenic progenitors (FAPs; Lin^−^/VCAM1^−^/Sca-1^+^), and double-negative cells (DN; Lin^−^/VCAM1^−^/Sca-1^−^), which harbor the Schwann cell population (Figure 2—figure supplement 1B). Flow cytometry analysis revealed that the vast majority of tdTomato-positive cells (∼92%) were located exclusively within the DN population, consistent with the expected identity of tSCs (Figure 2—figure supplement 1C-E). Finally, to confirm successful genetic recombination, we extracted genomic DNA from the sorted cell populations and performed PCR. The recombined tdTomato amplicon was strictly detected in the tdT+ fraction of the DN population, confirming that Cre-mediated recombination occurred specifically in the intended target cells (Figure 2—figure supplement 1F). These structural and genomic validations suggest that the *Col20a1-*CreERT2 mouse is a highly specific and robust tool for targeting tSCs in skeletal muscle.

Having confirmed the specificity, we next characterized the spatial distribution of the reporter expression across various skeletal muscles to establish the model as a universally reliable tool. We analyzed representative cranial (masseter, sternomastoid), axial (diaphragm, psoas), and appendicular (EDL, soleus) muscles, covering diverse anatomical locations and fiber type compositions. Immunofluorescence analysis revealed consistent and robust tdTomato labeling within tSCs across all examined muscle groups (Figure 2C). This uniform pattern confirms that the *Col20a1*-CreERT2 model can target tSCs globally throughout the body, regardless of their anatomical niche.

### Temporal mapping of *Col20a1*-CreERT2 expression during postnatal maturation

While the *Col20a1*-CreERT2 model showed high specificity in adult muscles, its application for other developmental stages requires a precise understanding of its expression dynamics. Thus, we investigated the temporal expression profile of this model during the critical window of NMJ maturation. *Col20a1*-tdT pups were administered a single intraperitoneal (IP) injection of tamoxifen (160 mg/kg body weight) at postnatal day 4 (P4), P7, P10, or P14, and analyzed at 4 weeks of age (Figure 3A). Interestingly, early induction at P4 and P7 resulted in tdTomato expression not only in tSCs but also partially in upstream axonal Schwann cells (aSCs) (Figure 3B, white arrowheads). However, this broader labeling became highly restricted as development progressed; induction at P10 and P14 yielded labeling that was largely confined to tSCs. These findings indicate that while *Col20a1* is broadly expressed in both tSCs and aSCs during early postnatal development, its expression becomes restricted to tSCs by P10. Consequently, we established P10 or later as the standard induction time point for selective tSC targeting.

**Figure 3.**
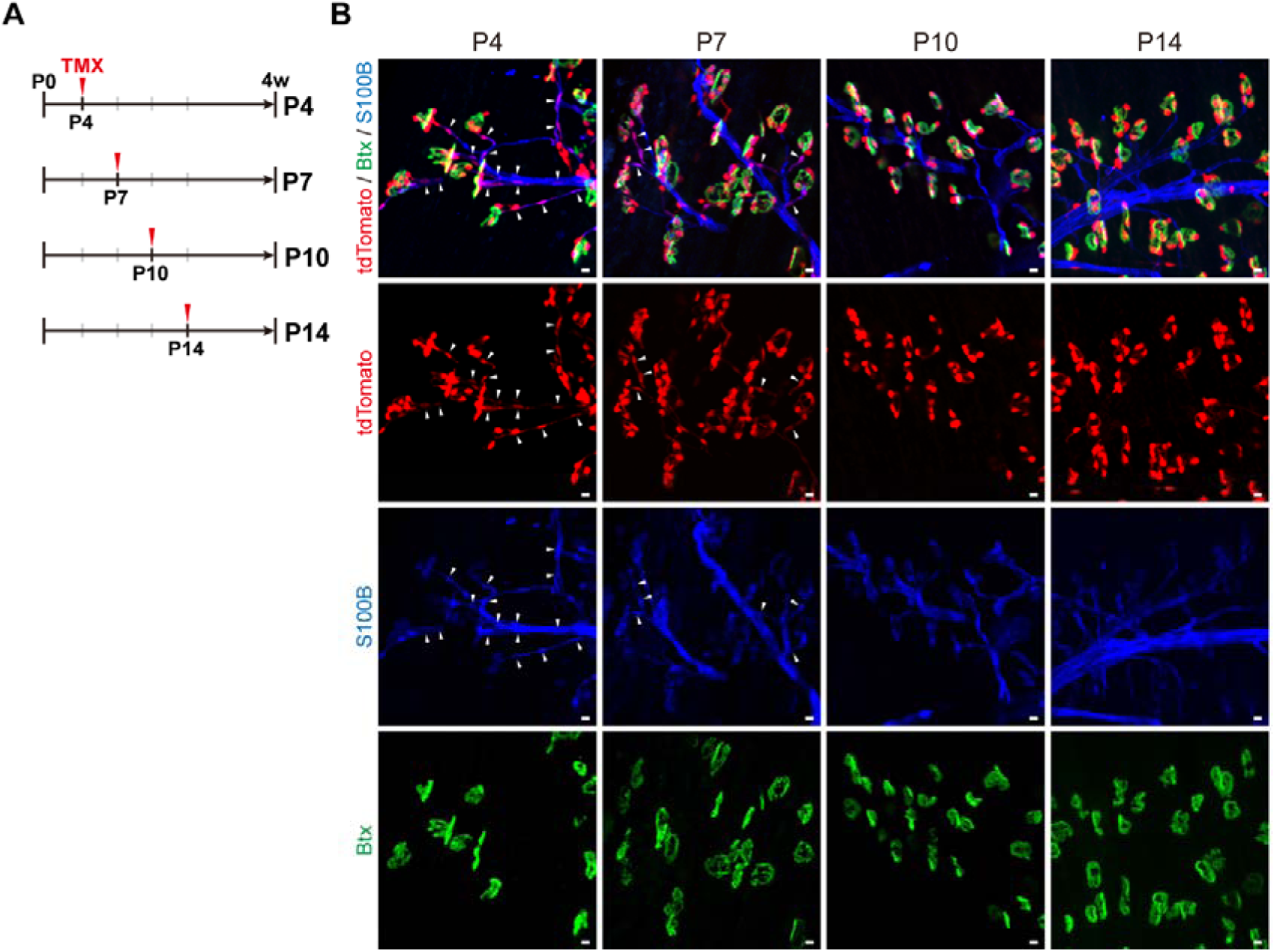
Temporal mapping establishes the postnatal specificity window of the *Col20a1*-CreERT2 model. (A) Experimental timeline for investigating the temporal dynamics of *Col20a1*-CreERT2 model. A single dose of tamoxifen (TMX, 160 mg/kg body weight) was administered in *Col20a1*-tdT mice via intraperitoneal injection at postnatal day 4 (P4), P7, P10, or P14, followed by analysis at 4 weeks (4w) of age. (B) Representative immunofluorescence images of EDL muscle NMJs corresponding to the temporal induction timeline. Early TMX induction at P4 and P7 results in tdTomato labeling (red) of both tSCs and portions of upstream axonal Schwann cells (aSCs, indicated by white arrowheads). In contrast, induction at P10 and P14 yields labeling that is largely confined to tSCs. S100B (blue); Btx (green). Scale bars: 10 µm.

### Targeted postnatal tSC ablation using the *Col20a1*-tdT/DTA model produces sustained reduction and subsequent recovery of glial coverage

Having established P10 as an induction time point for preferential tSC targeting (Figure 3), we sought to investigate the functional requirement of tSCs during postnatal NMJ maturation. We generated a genetic ablation model, *Col20a1*-CreERT2; *Rosa*-tdT/DTA (*Col20a1*-tdT/DTA), in which tamoxifen induction drives expression of diphtheria toxin subunit A (DTA) to ablate the targeted cells and of tdTomato to identify residual reporter-positive tSCs.

Previous work in adult mice has shown that glial coverage can be restored following tSC ablation (Hastings et al., 2020). We therefore mapped the time course of coverage loss and recovery to identify a suitable interval for phenotypic analysis. Following a single intraperitoneal injection of tamoxifen at P10, we examined NMJs in the EDL muscle at P14, P17, P21, and 4, 6, 8 and 12 weeks of age. We scored tdTomato labeling and S100B immunostaining separately, so that glial coverage could be assessed independently of reporter expression. Each NMJ was assigned to one of four categories: tdTomato-positive (tdT^+^)/S100B-positive (S100B^+^), tdT-negative (tdT^−^)/S100B-absent, tdT^−^/S100B-partial, and tdT^−^/S100B-full (Figure 4A, B). In control *Col20a1*-tdT mice examined in parallel, essentially all NMJs carried tdT^+^/S100B^+^ cells (Figure 4—figure supplement 1).

**Figure 4.**
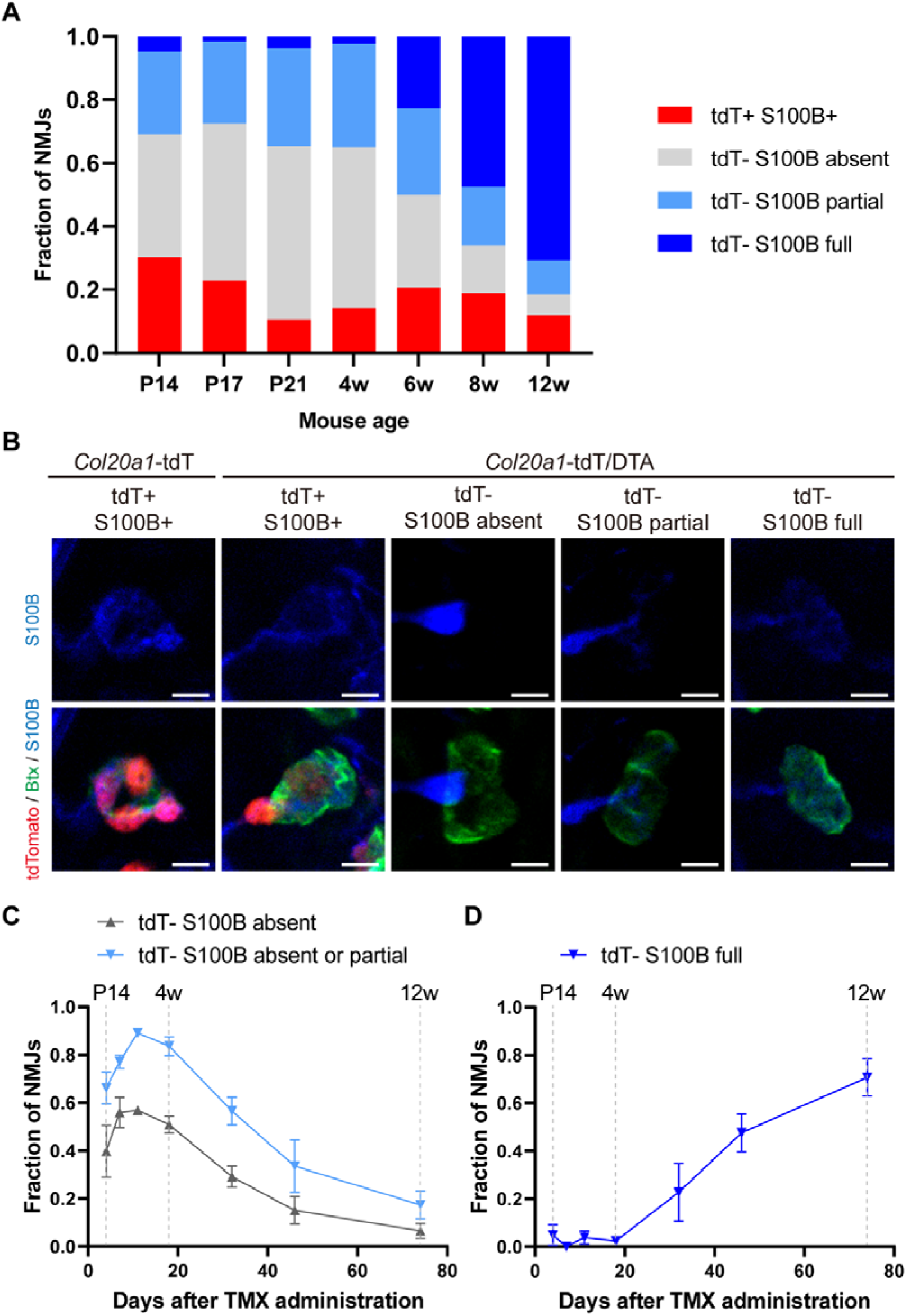
Postnatal tSC ablation produces a sustained loss of glial coverage followed by gradual recovery. Tamoxifen (TMX, 160 mg/kg body weight) was administered at postnatal day 10 (P10) via intraperitoneal injection, and NMJs of the EDL muscle were analyzed at P14, P17, P21 and 4, 6, 8 and 12 weeks of age. (A) Fraction of NMJs assigned to each of four categories at each time point: reporter-positive NMJs carrying S100B-positive tSCs (tdT^+^ S100B^+^), and reporter-negative NMJs in which S100B labeling was absent from the synaptic area (tdT^−^ S100B absent), covered it only in part (tdT^−^ S100B partial), or covered it completely as in controls (tdT^−^ S100B full). Glial coverage disruption is already evident at P14, when most NMJs show absent or partial S100B labeling, and NMJs with full coverage become progressively more frequent after 4 weeks of age. (B) Representative confocal images of each category, taken at P14. The leftmost column shows a control NMJ from a *Col20a1*-tdT mouse; the remaining columns show NMJs from *Col20a1*-tdT/DTA mice. Top row, S100B (blue); bottom row, merged tdTomato (red), α-bungarotoxin (Btx, green) and S100B (blue). Scale bars: 10 µm. (C, D) Fraction of NMJs in selected categories, plotted against time after tamoxifen administration; grey dashed lines mark the corresponding mouse ages. (C) NMJs lacking S100B coverage (tdT^−^ S100B absent), and the combined fraction of NMJs with absent or partial coverage. Both measures peak by P21 and decline thereafter, indicating that the loss of glial coverage is sustained but reversible. (D) NMJs with full S100B coverage in the absence of the tdTomato reporter (tdT^−^ S100B full). This population remains minimal through 4 weeks and then increases steadily, reaching approximately 71% of all NMJs by 12 weeks. The proportion of NMJs in each category was determined separately for each animal from all NMJs scored in that animal, and these per-animal proportions were then averaged. Data are presented as mean ± SD of n = 3 mice per time point (3,087 NMJs scored in total, on average 147 per mouse). The analysis characterizes the time course of coverage loss and recovery rather than testing specific contrasts; no statistical comparisons were performed.

At the earliest time point examined, P14, ∼30% of NMJs retained tdT+/S100B+ cells, whereas ∼65% showed absent or partial S100B coverage (Figure 4A, C). Glial coverage was therefore already disrupted by P14. The fraction of NMJs with absent or partial coverage peaked at P21 (∼86%) and remained high at 4 weeks (∼84%). It then declined as tdT^−^ NMJs with full S100B coverage became more frequent, reaching ∼71% of all NMJs by 12 weeks (Figure 4A, C, D).

Postnatal NMJ maturation, including the reorganization of postsynaptic AChR clusters and the development of junctional folds, continues through the first three postnatal weeks (Kariya et al., 2014; Medina-Moreno & Henríquez, 2022; Sanes & Lichtman, 1999, 2001). The coverage deficits in our model were evident by P14 and persisted through P21, thus overlapping the later stages of this period. We therefore selected 4 weeks of age for the subsequent structural and functional analyses, an age beyond the major maturation period at which glial coverage remained markedly reduced.

### Gross NMJ architecture is largely preserved after tSC ablation

Having established 4 weeks of age as the principal time point for phenotypic analysis, we examined gross NMJ morphology in EDL muscles following tamoxifen administration at P10. Immunofluorescence images of *Col20a1*-tdT/DTA muscle showed widespread loss of the tdTomato signal (Figure 5A), and high-magnification views of the outlined regions identified individual tdT^−^ NMJs with no detectable S100B coverage (Figure 5B), the population characterized in Figure 4.

**Figure 5.**
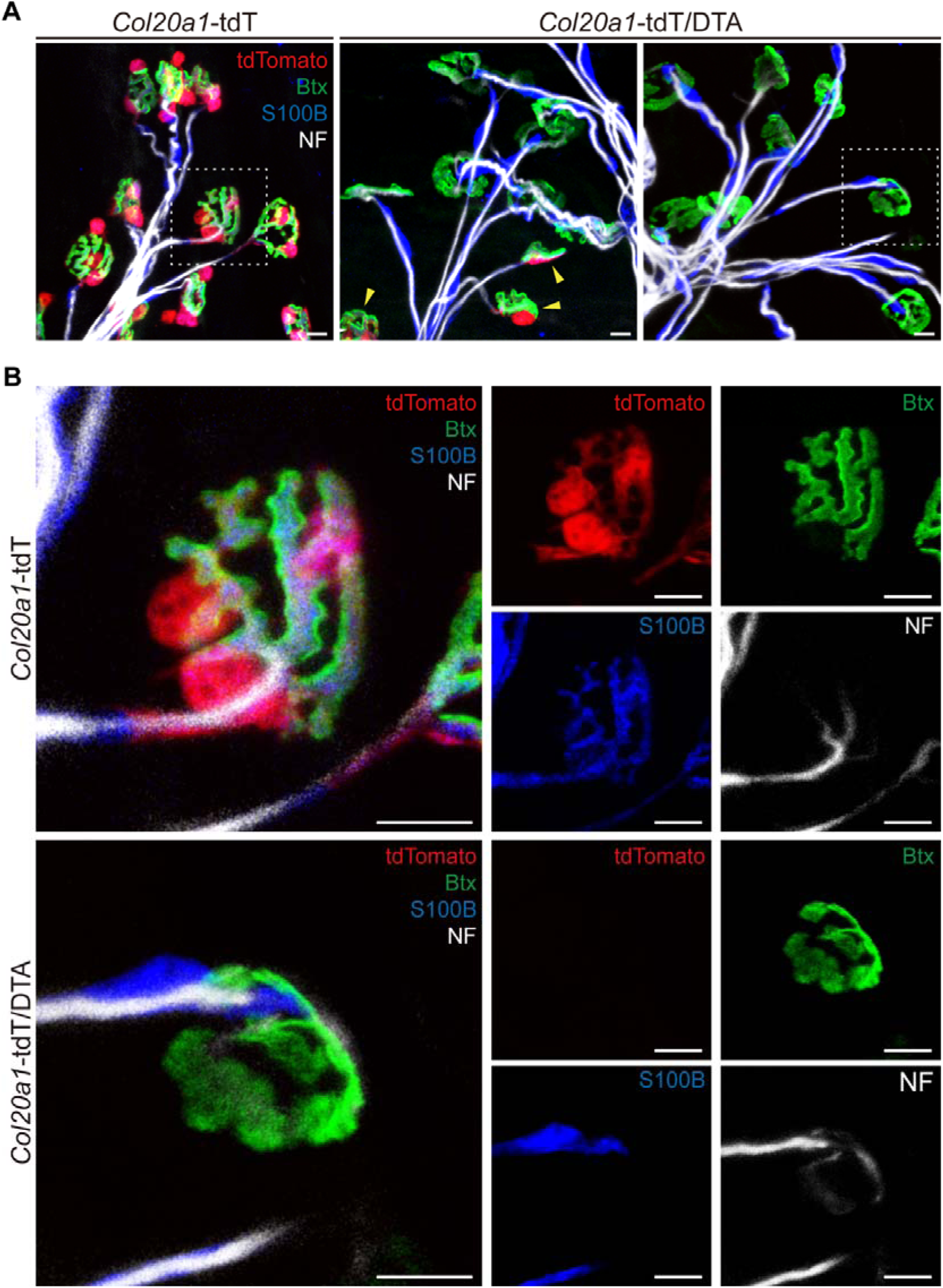
Gross NMJ architecture is largely preserved after tSC ablation. To induce tSC ablation, tamoxifen (TMX, 160 mg/kg body weight) was administered at postnatal day 10 (P10) via intraperitoneal injection, and NMJs were analyzed at 4 weeks of age. (A) Representative low-magnification immunofluorescence images of EDL muscle NMJs from *Col20a1*-tdT control (left) and *Col20a1*-tdT/DTA mice (middle and right), labeled for tdTomato (red), α-bungarotoxin (Btx, green), S100B (blue) and neurofilament (NF, white). Compared to controls, *Col20a1*-tdT/DTA muscles show a widespread loss of the tdTomato signal. Yellow arrowheads indicate rare surviving “escaper” tSCs, and white dashed boxes outline the regions shown at higher magnification in B. Scale bars: 10 µm. (B) High-magnification confocal images of individual NMJs from *Col20a1*-tdT control (top) and tSC-ablated *Col20a1*-tdT/DTA mice (bottom), shown as merged images (left) and as single channels (right). In *Col20a1*-tdT/DTA mice, NMJs lacking both the tdTomato signal and S100B coverage retain single-axon innervation with no overt signs of denervation or axon terminal pathology, and their AChR clusters show neither severe fragmentation nor morphological immaturity. Scale bars: 10 µm.

Unexpectedly, despite these pronounced coverage deficits, the overall architecture of both the presynaptic and postsynaptic compartments remained largely intact. Previous work in developing amphibian NMJs showed that loss of perisynaptic Schwann cells can lead to widespread nerve terminal retraction (Reddy et al., 2003). In our model, however, NMJs displayed single-axon innervation without overt signs of denervation or axon terminal pathology at 4 weeks (Figure 5B).

Postsynaptic AChR clusters also undergo extensive structural remodeling during the first three postnatal weeks (Kariya et al., 2014; Marques et al., 2000). Although the coverage deficits overlapped this period, AChR clusters did not exhibit severe fragmentation or morphological immaturity (Figure 5B). Collectively, these observations indicate that gross presynaptic and postsynaptic architecture can remain largely preserved despite substantial disruption of tSC coverage during postnatal maturation.

### Systemic ablation of tSCs does not impair overall growth or gross motor behavior

As our newly developed model offers the advantage of system-wide tSC targeting, it enables the assessment of whole-body physiological impacts. Thus, we next investigated whether widespread tSC loss translates to macroscopic physiological or behavioral deficits. We first monitored general somatic growth of tSC-ablated mice by measuring body weight at the time of tamoxifen induction (P10) and post-ablation at 4 weeks of age. We observed no significant differences in body weight between *Col20a1*-tdT controls and *Col20a1*-tdT/DTA mice at either time point, indicating normal growth trajectories (Figure 6A, B). To assess overall motor coordination and muscle function, we performed a beam walking test (Lukong & Richard, 2008; Luong et al., 2011; Quinn et al., 2007). The latency to cross the beam was comparable between the ablated mice and controls, indicating normal balance and gross motor coordination (Figure 6C). Furthermore, we conducted a gait analysis to evaluate fine motor control, a metric that is often altered in classical neuromuscular disorders (Maricelli et al., 2016; Preisig et al., 2016; Sheppard et al., 2020; Wooley et al., 2005). Consistent with the beam walking results, systemic tSC ablation did not significantly alter either stride length or stride width (Figure 6D, E). Taken together with our morphological data (Figure 4), these behavioral results indicate that tSCs are dispensable for overall somatic growth and basic motor behavior during the postnatal maturation period. This lack of a severe macroscopic phenotype suggests that tSCs do not govern broad neuromuscular integrity.

**Figure 6.**
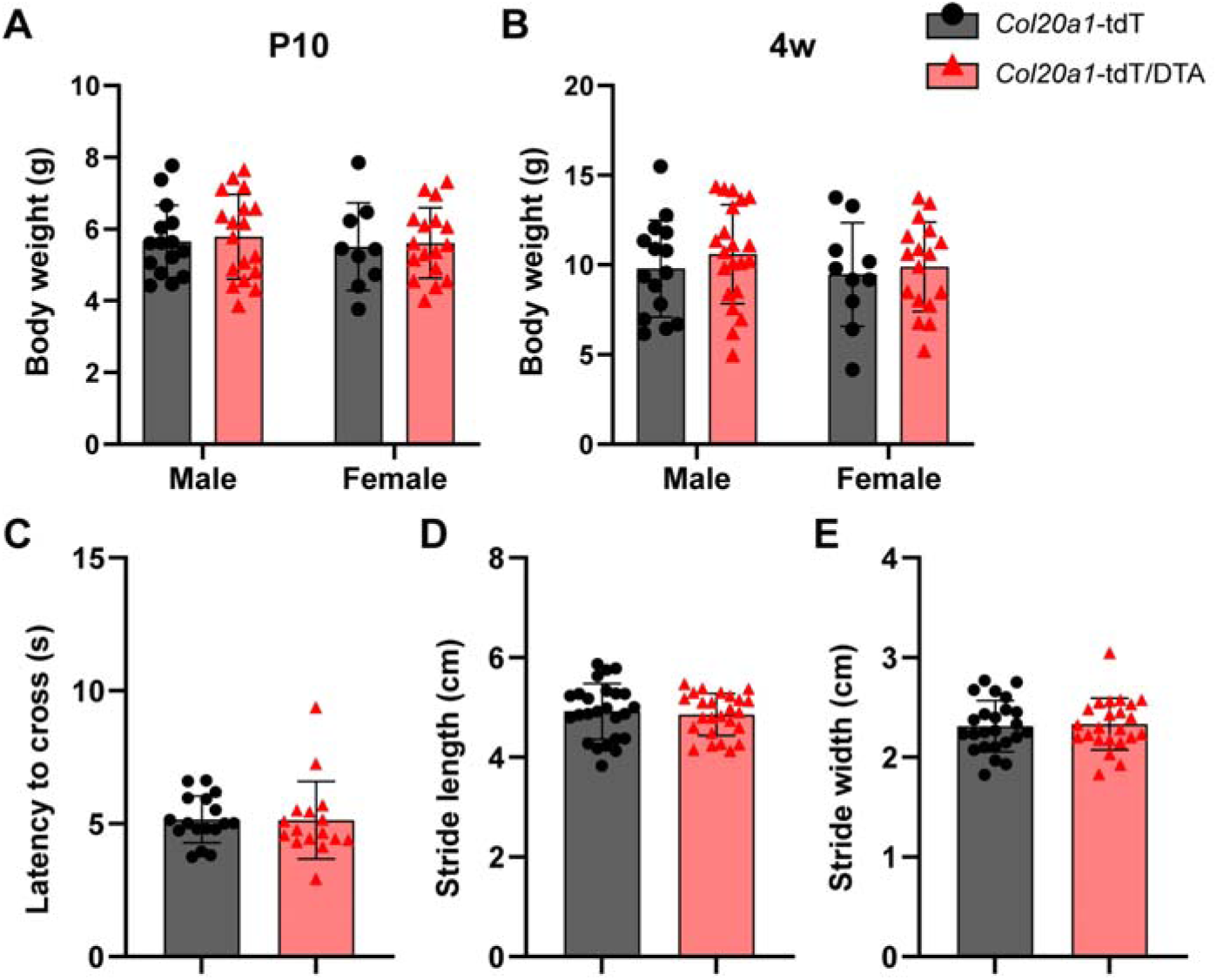
Systemic ablation of tSCs does not impair overall somatic growth or gross motor behavior. (A, B) Body weight analysis of male and female control (*Col20a1*-tdT) and tSC-ablated (*Col20a1*-tdT/DTA) mice at (A) postnatal day 10 (P10, time of induction; control: n = 15 males, 9 females; ablated: n = 18 males, 17 females) and (B) 4 weeks of age (4w, post-ablation; control: n = 15 males, 10 females; ablated: n = 21 males, 17 females). Systemic loss of tSCs does not significantly alter normal growth trajectories in either sex. (C) Assessment of gross motor coordination using the beam walking test. The latency to cross the beam is comparable between the ablated mice and controls at 4 weeks of age (control, n = 17 mice; ablated, n = 16 mice). (D, E) Quantitative gait analysis evaluating fine motor control. Systemic tSC ablation does not significantly affect (D) stride length or (E) stride width at 4 weeks of age (n = 25 mice per group). Data are presented as mean ± SD. Statistical significance was determined using two-way ANOVA with Šídák’s multiple comparisons test for A and B, and Student’s t-test for C, D, and E. All the comparisons did not reach statistical significance (*p* > 0.05).

### tSC ablation impairs presynaptic vesicle homeostasis and high-frequency synaptic transmission

To determine whether tSCs are required for fine-tuned synaptic function despite normal morphology, we performed electrophysiological recordings. We first measured miniature endplate potentials (mEPPs) (Figure 7A) and found that mutant NMJs exhibited a significant increase in mEPP frequency compared to controls (Figure 7D, E), while mEPP amplitude was significantly decreased (Figure 7B, C). These results indicate an increased frequency of spontaneous ACh release and a smaller postsynaptic response to individual release events in the absence of tSCs.

**Figure 7.**
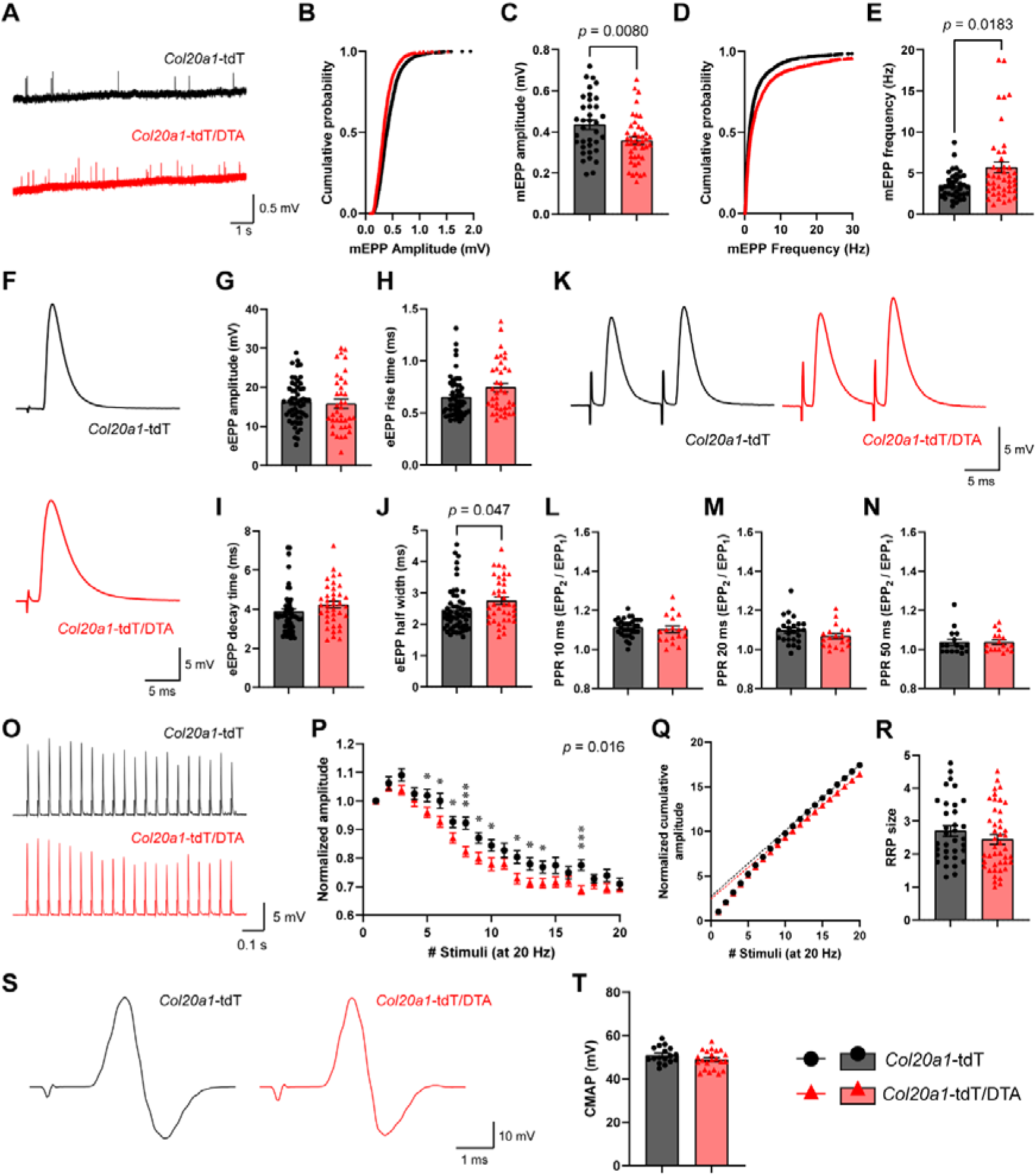
Targeted tSC ablation impairs presynaptic vesicle homeostasis and compromises high-frequency synaptic transmission. Tamoxifen (TMX, 160 mg/kg body weight) was administered at postnatal day 10 (P10) via intraperitoneal injection, and electrophysiological recordings were performed at 4 weeks of age. Black, *Col20a1*-tdT (control); red, *Col20a1*-tdT/DTA (tSC-ablated). (A) Representative electrophysiological traces of miniature endplate potentials (mEPPs) recorded from isolated EDL muscles. (B, C) Cumulative probability distribution (B) and mean (C) of mEPP amplitude, which is reduced in *Col20a1*-tdT/DTA mice, indicating a smaller postsynaptic response to individual release events. (D, E) Cumulative probability distribution (D) and mean (E) of mEPP frequency, which is increased in *Col20a1*-tdT/DTA mice, indicating an increased spontaneous ACh release. (F) Representative traces of single evoked endplate potentials (eEPPs). (G–J) Quantification of eEPP kinetics. eEPP amplitude (G), rise time (H), decay time (I) and half width (J). Mutant NMJs exhibit significantly increased eEPP half width (J). There are no significant differences in amplitude (G), rise time (H) or decay time (I). (K) Representative traces of paired-pulse responses at an inter-stimulus interval of 10 ms. (L–N) Paired-pulse ratio (PPR, EPP_2_/EPP_1_) at 10 ms (L), 20 ms (M) and 50 ms (N) inter-stimulus intervals, which is comparable between groups, indicating a preserved basal release probability in the absence of tSCs. (O) Representative eEPP trains elicited by repetitive 20 Hz stimulation (20 pulses). (P) Normalized eEPP amplitudes during the 20 Hz stimulation train. *Col20a1*-tdT/DTA mice NMJs exhibit a progressively more severe synaptic depression compared to controls. (Q, R) Cumulative amplitude plot (Q) and estimation of the readily releasable pool (RRP) size (R) calculated via back-extrapolation of the cumulative eEPP amplitude. Calculated RRP size is not significantly altered between groups. (S) Representative traces of Compound Muscle Action Potential (CMAP) recorded from the gastrocnemius. (T) Quantification of CMAP amplitude, indicating normal macroscopic physiological function in mutant mice, consistent with behavioral data. Data are presented as mean ± SEM with individual values shown. mEPP (A–E): control, n = 36 NMJs from 6 mice; tSC-ablated, n = 43 NMJs from 8 mice. eEPP (F–J): control, n = 55 NMJs from 13 mice; tSC-ablated, n = 38 NMJs from 10 mice. PPR (K–N): at 10 ms, control n = 33 NMJs from 9 mice and tSC-ablated n = 18 NMJs from 6 mice; at 20 ms, control n = 25 NMJs from 7 mice and tSC-ablated n = 19 NMJs from 6 mice; at 50 ms, control n = 16 NMJs from 4 mice and tSC-ablated n = 17 NMJs from 6 mice. Train and RRP (O–R): control, n = 34 NMJs from 8 mice; tSC-ablated, n = 44 NMJs from 10 mice. CMAP (S, T): control, n = 17 mice; tSC-ablated, n = 22 mice. Statistical significance was determined using Student’s t-test for C, G–J, L–N, R, and T, the Mann-Whitney U test for E, and Repeated Measures ANOVA with Bonferroni’s multiple comparisons for P. Exact *p*-values are shown for comparisons that reached statistical significance (*p* < 0.05). For P, the overall effect of genotype is given and asterisks indicate individual stimulus numbers (\**p* < 0.05, \*\**p* < 0.01, \*\*\**p* < 0.001).

Analysis of single evoked endplate potentials (eEPPs) (Figure 7F) revealed no significant difference in amplitude between mutant and control NMJs (Figure 7G). However, eEPP half width was significantly increased in mutants (Figure 7J), whereas rise time and decay time did not differ significantly between groups (Figure 7H, I). The paired-pulse ratio (PPR) was comparable between groups across all intervals (Figure 7K–N). These findings suggest that the strength of single-stimulus responses and initial release probability were largely preserved following tSC ablation.

Despite the preservation of single-stimulus eEPP amplitude, the functional endurance of mutant NMJs was compromised under high-demand conditions. Specifically, under repetitive 20 Hz stimulation (eEPP-train), mutant NMJs exhibited significantly greater depression of normalized eEPP amplitudes compared to controls (Figure 7O, P). Estimation of the readily releasable pool (RRP) size via back-extrapolation of cumulative amplitude revealed no significant difference between the two groups (Figure 7Q, R). Finally, CMAP amplitude did not differ significantly between groups (Figure 7S, T), consistent with the comparable single-stimulus eEPP amplitudes and the lack of gross behavioral deficits. Collectively, these results indicate that tSCs are essential for maintaining presynaptic vesicle homeostasis and sustaining high-frequency synaptic transmission.

### Ultrastructural analysis reveals reduced presynaptic vesicle density following tSC ablation

To examine the ultrastructural changes accompanying the electrophysiological alterations, we performed electron microscopy (EM) on EDL muscle NMJs at 4 weeks of age (Figure 8A). NMJs were classified into three groups: *Col20a1*-tdT controls, *Col20a1*-tdT/DTA NMJs with retained tSCs (tSC-retained), and *Col20a1*-tdT/DTA NMJs lacking identifiable tSC coverage (tSC-ablated).

**Figure 8.**
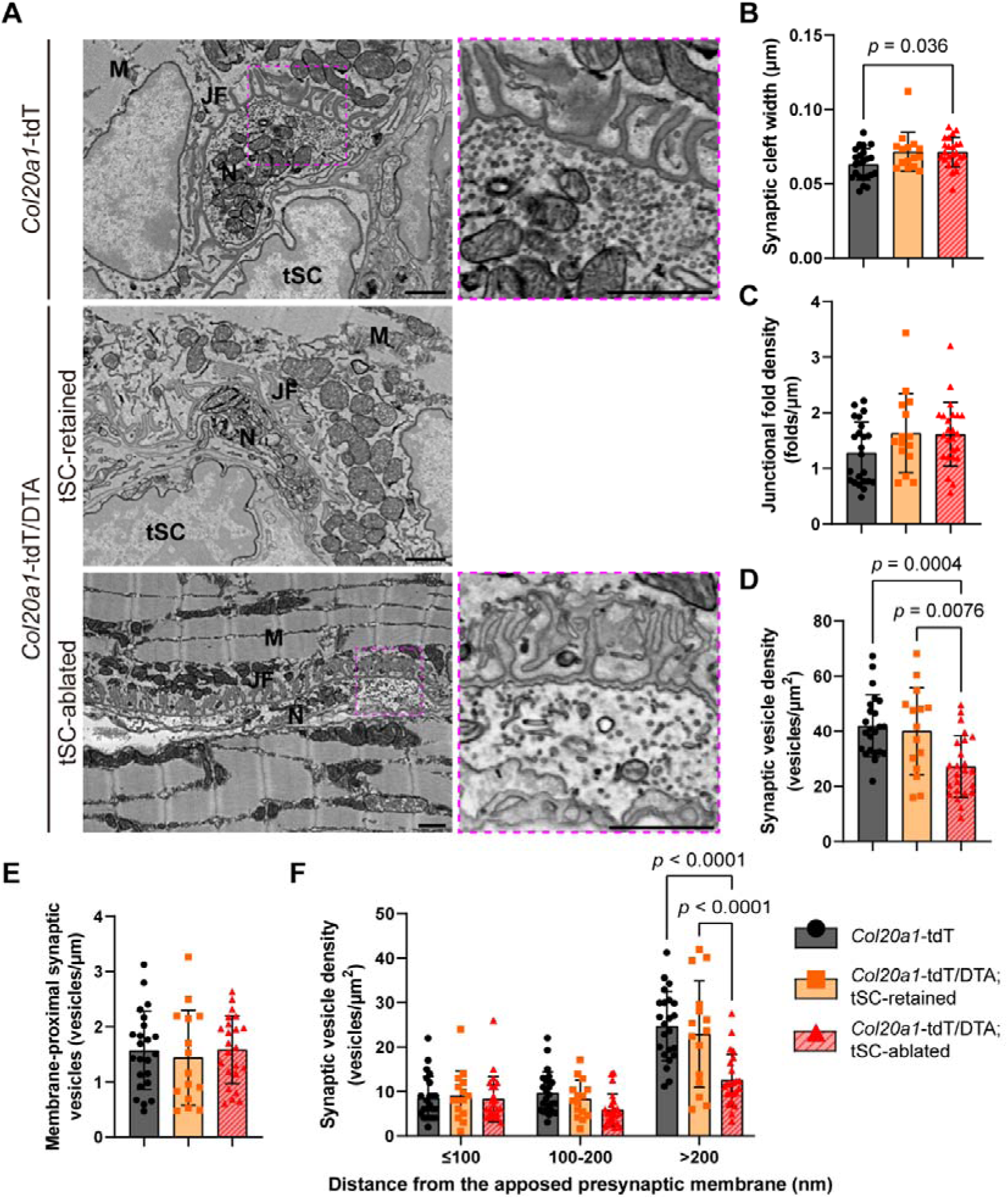
Ultrastructural analysis reveals a position-dependent reduction in synaptic vesicle density following tSC ablation. Tamoxifen (TMX, 160 mg/kg body weight) was administered at postnatal day 10 (P10) via intraperitoneal injection, and NMJs of the EDL muscle were analyzed at 4 weeks of age. (A) Representative backscattered scanning electron microscopy (BSE-SEM) micrographs of ultra-thin sections of NMJs from control (*Col20a1*-tdT), mutant NMJs with retained tSCs (*Col20a1*-tdT/DTA; tSC-retained), and mutant NMJs with ablated tSCs (*Col20a1*-tdT/DTA; tSC-ablated). High-magnification views of the regions indicated by pink dashed boxes are shown on the right. M, muscle; N, nerve terminal; JF, junctional folds; tSC, terminal Schwann cell. Scale bars: 1 µm. (B, C) Quantitative analysis of (B) synaptic cleft width and (C) junctional fold density. Synaptic cleft width is modestly increased in tSC-ablated NMJs relative to controls, whereas junctional fold density is unchanged across all groups. (D) Quantification of total synaptic vesicle density. The tSC-ablated NMJs exhibit a significant reduction in vesicle density compared to both control and tSC-retained groups. (E) Density of membrane-proximal synaptic vesicles, defined as vesicles whose center lies within 50 nm of the apposed presynaptic membrane, measured perpendicular to that membrane and normalized to its length. This population is comparable across all groups. (F) Synaptic vesicle density binned by distance from the apposed presynaptic membrane (≤ 100, 100−200 and >200 nm). The reduction in tSC-ablated NMJs is confined to the >200 nm bin. Data are presented as mean ± SD (control, n = 23 NMJs from 6 mice; mutant NMJs with retained tSCs, n = 15 NMJs from 6 mice; mutant NMJs with ablated tSCs, n = 23 NMJs from 5 mice). Statistical significance was determined using one-way ANOVA with Bonferroni’s post hoc test for B– E and two-way ANOVA with Šídák’s multiple comparisons test for F. Exact *p*-values are shown for comparisons that reached statistical significance (*p* < 0.05).

Synaptic cleft width was modestly but significantly increased in tSC-ablated NMJs compared with controls (Figure 8B), whereas no significant differences in junctional fold density were detected among the three groups (Figure 8C). Thus, although gross NMJ architecture appeared largely preserved, tSC ablation was accompanied by a subtle alteration in synaptic cleft geometry.

Notably, synaptic vesicle density was significantly reduced in tSC-ablated NMJs compared with both control and tSC-retained NMJs (Figure 8D). In the preceding electrophysiological analysis, however, train-derived RRP estimates were broadly similar between groups. We therefore examined whether this functional observation was reflected in the ultrastructural distribution of synaptic vesicles. As a complementary anatomical measure, we quantified membrane-proximal vesicles within 50 nm of the apposed presynaptic membrane, a distance approximating the diameter of a single synaptic vesicle. Their number, normalized to the presynaptic apposition length, did not differ significantly among the three groups (Figure 8E). Thus, the reduction in overall vesicle density was not accompanied by a detectable decrease in membrane-proximal vesicles, a pattern compatible with the broadly similar electrophysiological RRP estimates.

Given the reduction in overall vesicle density without a detectable change in the membrane-proximal measure, we next examined whether the reduction was more evident farther from the apposed presynaptic membrane. Vesicles were classified into distance bins, and counts in each bin were normalized to the total terminal profile area. Vesicle density in the >200 nm bin was significantly lower in tSC-ablated NMJs than in both control and tSC-retained NMJs, whereas no significant differences were detected in the ≤100 nm or 100–200 nm bins (Figure 8F). Thus, the overall reduction in vesicle density was most apparent in the distal vesicle population.

Overall, the reduced presynaptic vesicle density helps explain the more pronounced depression during repetitive stimulation, even though single-stimulus eEPP amplitudes were largely maintained. Together with increased spontaneous release, these findings are consistent with impaired presynaptic vesicle homeostasis after tSC ablation, potentially accelerating vesicle depletion during repeated activity. Thus, tSCs appear to contribute to presynaptic vesicle homeostasis, with a possible additional role in maintaining synaptic cleft geometry.

### Long-term analysis reveals preserved neuromuscular function following postnatal tSC ablation

At 4 weeks of age, we found loss of tSCs induced changes in vesicle homeostasis and ultrastructure while leaving gross NMJ morphology and behavioral phenotype intact. Our time-course analysis showed that incomplete glial coverage persisted for several weeks after P10 induction, followed by a gradual increase in full S100B coverage toward 12 weeks of age (Figure 4). We therefore examined NMJ morphology and neuromuscular function at 12 weeks to assess whether they remained preserved during this later stage of recovery.

Consistent with the recovery of glial coverage reported after tSC ablation in adult mice (Hastings et al., 2020), immunofluorescence analysis identified NMJs with extensive S100B coverage but no detectable tdTomato signal in *Col20a1*-tdT/DTA mice (Figure 9A, B). These NMJs retained AChR clusters associated with presynaptic terminals, with no overt signs of denervation or severe fragmentation. Thus, the recovery of glial coverage was accompanied by largely preserved gross NMJ morphology.

**Figure 9.**
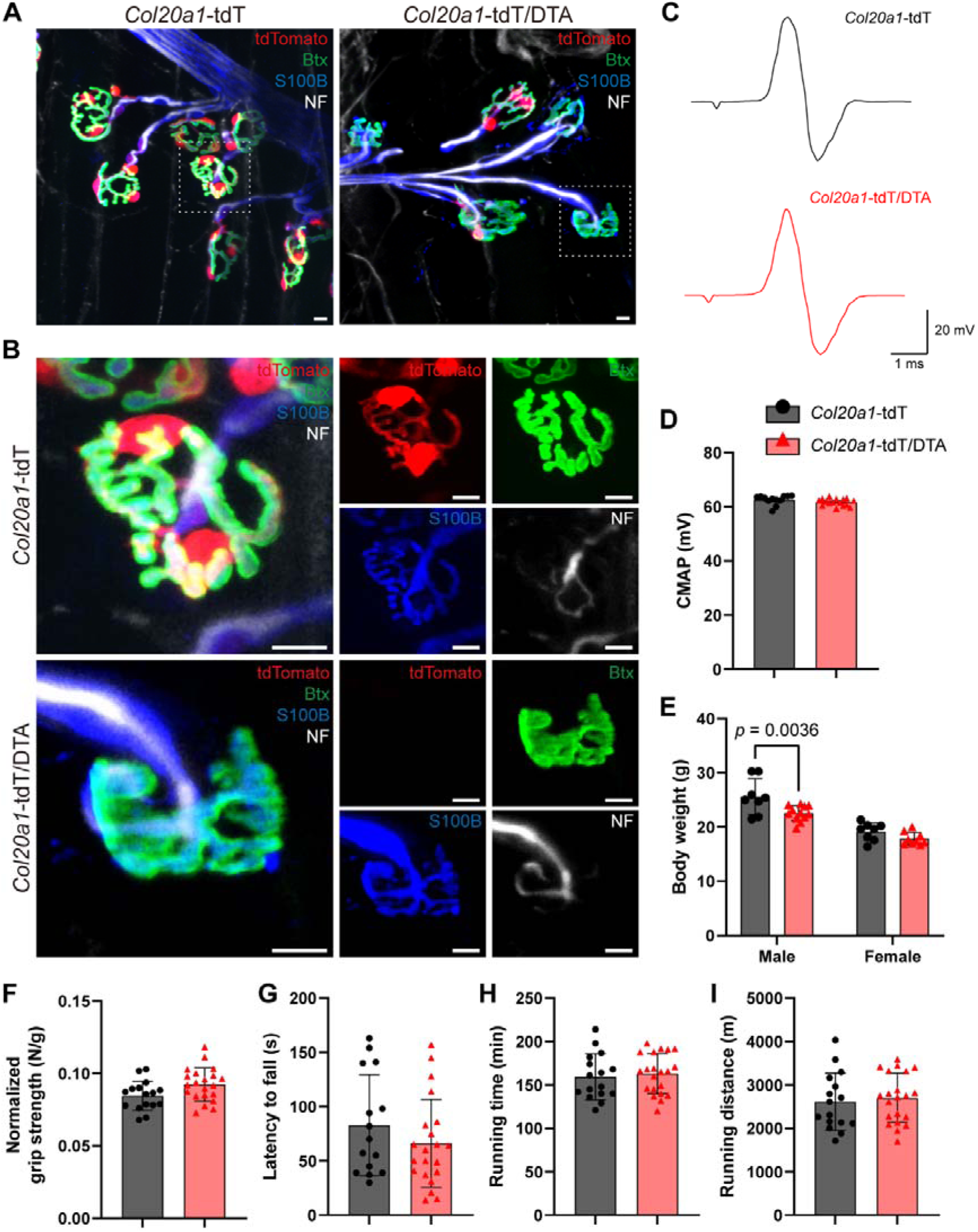
Long-term repopulation of Schwann cells maintains structural integrity and gross neuromuscular function. To assess long-term recovery, tamoxifen (TMX, 160 mg/kg body weight) was administered at postnatal day 10 (P10) by intraperitoneal injection, and mice were analyzed at 12 weeks of age. (A) Representative immunofluorescence images of muscle NMJs from *Col20a1*-tdT control (left) and tSC-ablated *Col20a1*-tdT/DTA mice (right) at 12 weeks of age. White dashed boxes indicate regions magnified in B. Scale bars: 10 µm. (B) High-magnification confocal images of individual NMJs from *Col20a1*-tdT (top) and *Col20a1*-tdT/DTA (bottom) mice. In the mutant NMJs, S100B^+^ glial cells (blue) repopulate the synapse despite the persistent absence of the tdTomato reporter (red). The gross architecture of presynaptic motor axons (NF, white) and postsynaptic AChR clusters (Btx, green) is preserved. Scale bars: 10 µm. (C) Representative traces of Compound Muscle Action Potential (CMAP). (D) Quantification of CMAP amplitude, indicating sustained physiological function in tSC-ablated mice (control, n = 12 mice; ablated, n = 16 mice). Data are presented as mean ± SEM. (E) Body weight analysis of male and female mice at 12 weeks of age (control: n = 8 males, 8 females; ablated: n = 12 males, 9 females). Data are presented as mean ± SD. (F–I) Macroscopic motor behavioral assessments showing normal long-term function in mutant mice, including (F) normalized grip strength (control, n = 16 mice; ablated, n = 21 mice), (G) motor coordination assessed by latency to fall in the rotarod test (control, n = 15 mice; ablated, n = 21 mice), and endurance running capacity measured by (H) running time and (I) running distance (control, n = 16 mice; ablated, n = 21 mice). Data are presented as mean ± SD. Statistical significance was determined using two-way ANOVA with Šídák’s multiple comparisons test for E and Student’s t-test for D and F–I. Exact *p*-values are shown for comparisons that reached statistical significance (*p* < 0.05).

Furthermore, we assessed various functional and behavioral metrics to confirm that this repopulation successfully supports long-term neuromuscular health. Similar to our observations at 4 weeks, we found no significant differences in CMAP amplitudes (Figure 9C, D). Regarding gross physiological indicators, while a slight reduction in body weight was observed in tSC-ablated males at 12 weeks (Figure 9E), this did not translate into any macroscopic motor impairment. Indeed, all motor behavioral assessments, including grip strength (Figure 9F), rotarod motor coordination (Figure 9G), and endurance running capacity (Figure 9H, I), remained comparable to controls. Together, these data suggest that the neuromuscular system possesses a robust compensatory mechanism to replace lost tSCs, thereby maintaining overall synaptic stability and sustaining fundamental motor function.

## Discussion

In this study, we identified *Col20a1* in a transcriptomic screen as a gene highly enriched in tSCs and established a *Col20a1*-CreERT2 mouse model to investigate their functional roles *in vivo*. Targeted tSC ablation from P10 produced a sustained loss of glial coverage that was already evident by P14 and persisted through 4 weeks of age. Over this period, gross NMJ architecture and motor behavior were preserved, whereas miniature endplate potentials became more frequent and smaller in amplitude, depression during repetitive stimulation was accelerated, and synaptic vesicle density was reduced, with the loss confined to vesicles lying distant from the presynaptic membrane. Glial coverage was subsequently restored by Schwann cells that had not undergone recombination, underscoring the dynamic plasticity of the tripartite synapse.

A long-standing challenge in tSC biology has been the difficulty of manipulating these cells without also affecting other Schwann cell lineages (Hastings & Valdez, 2024; Jablonka□Shariff et al., 2021; Santosa et al., 2018). Much of the available genetic access has come from pan-Schwann cell lines, such as those based on *Plp1* and *S100b* (Barik et al., 2016; Zuo et al., 2004), in which tSCs are labeled or manipulated together with the myelinating Schwann cells that ensheathe motor axons. Effects attributable to tSCs are therefore difficult to separate from those of concurrent aSC manipulation (Barik et al., 2016; Doerflinger et al., 2003). Acute physiological approaches such as calcium imaging offer high resolution (Darabid et al., 2013; Reist & Smith, 1992; Smith et al., 2013) but are suited to localized observation rather than long-term, organism-wide assessment. More selective genetic tools have since narrowed this gap. Combined S100b and NG2 reporters allow synaptic Schwann cells to be identified and isolated (Castro et al., 2020). An inducible Kir4.1-CreERT2 line provides access to non-myelinating Schwann cells, including tSCs (Procacci et al., 2023). More recently, *Col20a1* has been deleted in Schwann cells and globally to examine the function of the protein (Hastings et al., 2025). What has remained unavailable is an inducible driver whose recombination is largely confined to tSCs within muscle, permitting their selective removal and the assessment of consequences over time in the intact animal.

The *Col20a1-*CreERT2 model provides this access. Tamoxifen induction at P10 confined recombination largely to tSCs and allowed the consequences of their ablation to be followed through 12 weeks of age. A small fraction of tdTomato-positive cells fell outside the sorted population containing tSCs (Figure 2—figure supplement 1), indicating that some other cell types may also be labeled. The pattern nonetheless differs from that of pan-Schwann cell lines. tdTomato did not extend along motor axons in the chain-like arrangement characteristic of myelinating Schwann cells, whose concurrent manipulation is the principal confound of *Plp1*- and *S100b*-based tools. Because the ablation is genetic and applied systemically, its consequences could also be followed in the intact animal, from synaptic ultrastructure to motor behavior, over a timescale that acute preparations do not reach.

The labeling pattern also changed over development. At P4 and P7, tdTomato was present in a few preterminal Schwann cells along motor axons in addition to tSCs, and this component was no longer evident with induction from P10 onward (Figure 3). This may reflect a property of the marker itself. *Col20a1* could be expressed early in both preterminal Schwann cells and the cells that later form tSCs, or in a progenitor common to both, with expression becoming confined to tSCs over the second postnatal week. Alternatively, the cells labeled along axons at these early stages may themselves migrate to the synapse and join the tSC population. However, our induction experiment reports where recombination occurred rather than the fate of the recombined cells and cannot separate the two accounts. Following the fate of the cells labeled at these earlier stages would be an interesting use of this model. Lineage tracing could establish whether they migrate to the synapse and contribute to the tSC population.

Conventional wisdom, supported by several experimental models, holds that tSCs are required for the structural maintenance of the NMJ. Antibody-mediated ablation of perisynaptic Schwann cells at frog NMJs led to nerve terminal retraction within a week (Koirala et al., 2003). Mice in which Schwann cells were targeted with *Plp1*-CreERT2 showed both pre- and postsynaptic structural collapse (Barik et al., 2016). Acute application of patient-derived sera to isolated muscle produced structural abnormalities within an hour (Jablonka-Shariff et al., 2023). Together these findings established the view that loss of tSCs triggers structural failure at the synapse. In this study, however, the outcome of sustained tSC ablation was unexpected. NMJs showed neither denervation nor AChR fragmentation, even though glial coverage was already lost by P14 and remained substantially reduced at the time of analysis (Figure 4).

These experiments differ in ways that may account for the discrepancy. In models based on pan-glial promoters, tSCs are removed together with the Schwann cells that support the motor axon, so the structural collapse reported may reflect this combined loss rather than the loss of tSCs alone (Barik et al., 2016). In antibody-based approaches, the complement response raised against tSCs may also extend to the off-target nerve terminal (Halstead et al., 2004). DTA, by contrast, acts cell-autonomously and induces apoptosis in the cells that express it, so the effects of ablation are largely confined to those cells. By confining the perturbation largely to tSCs, the present model reveals that the mammalian NMJ is structurally far more resilient to the loss of its glial partner than previously appreciated.

Beyond the maintenance of existing structures, our model also addresses whether tSCs are required for the maturation of the NMJ. Several hallmarks of postnatal maturation, including the remodeling of AChR clusters into their mature “pretzel-like” topology and the development of junctional folds, continue through the third postnatal week (Kariya et al., 2014; Marques et al., 2000; Matthews-Bellinger & Salpeter, 1983; Shi et al., 2012). In our model, glial coverage was substantially reduced over the later part of this period. At 4 weeks, AChR clusters showed neither fragmentation nor an immature plaque-like morphology, and junctional fold density was unchanged, indicating that these aspects of maturation can proceed without normal tSC coverage. Earlier events are less accessible in this model. Induction at P10 was required for recombination to be largely restricted to tSCs (Figure 3), and the loss of glial coverage became evident at P14 (Figure 4), by which time synapse elimination is largely complete (Balice-Gordon & Lichtman, 1993; Duxson, 1982; Favero et al., 2015; Hirata et al., 1997; Lanuza et al., 2002). Whether tSCs are required for this process, in which they have been implicated (Lee et al., 2016), could not be addressed here, and their contribution at these earlier stages may well be greater than the present data can show.

Despite the preservation of macroscopic structure, our electrophysiological and ultrastructural analyses revealed defects in presynaptic homeostasis. An increased frequency of spontaneous ACh release and accelerated synaptic fatigue during high-frequency stimulation were accompanied by a reduction in synaptic vesicle density. Enhanced synaptic depression despite comparable RRP estimates raises the possibility that tSC ablation limits vesicle availability during repetitive stimulation, through increased resting release or impaired vesicle replenishment.

The ultrastructural loss was not uniform across the terminal, and its distribution bears on this. Vesicles lying close to the presynaptic membrane were as numerous as in controls, and the reduction was confined to those positioned further away. As vesicles already nearest to the membrane are the ones most likely to be available for immediate release, this distribution parallels what was observed physiologically: responses to single stimuli, including eEPP amplitude, were unchanged, and the deficit emerged only when release was sustained. Although these observations do not establish selective reduction of a functionally defined reserve pool, they indicate that tSCs are required for maintaining the presynaptic vesicle pool.

We showed that the spontaneous release was also changed in amplitude as well as frequency. The reduced mEPP amplitudes indicate smaller postsynaptic responses to individual release events, while comparable eEPP amplitudes may reflect compensatory maintenance of evoked transmission. Furthermore, increased eEPP half width despite unchanged rise and decay times suggests altered temporal regulation of evoked responses. EM analysis also revealed synaptic cleft widening following tSC ablation. This structural change may contribute to the reduced quantal responses and prolonged evoked responses, although a causal relationship remains to be established.

These findings suggest that tSCs function as active signaling modulators of the nerve terminal. Several mechanisms may underlie this functional regulation. For example, given that tSCs are known to sense nerve terminal activity via purinergic or calcium signaling (Darabid et al., 2013; Darabid et al., 2018; Heredia et al., 2018; Jahromi et al., 1992; Robitaille, 1998), they may provide a feedback loop that supports vesicle replenishment. Alternatively, tSCs may regulate the perisynaptic ionic environment by buffering excess potassium or calcium (Heredia et al., 2018; Mi et al., 1996); thus, their absence could leave the nerve terminal chronically more excitable, which would be consistent with the increase in spontaneous release. Further studies are required to decipher the exact molecular cross-talk responsible for this tSC-mediated vesicle homeostasis.

It is noteworthy that the electrophysiological alterations did not translate into overt motor deficits. The discrepancy is less marked than it may appear. Neuromuscular transmission operates with a considerable safety factor, and under low-frequency conditions the evoked response and the CMAP were unchanged, so ordinary locomotion need not be compromised. What was impaired was the supply of vesicles during sustained activity, and a deficit of this kind would emerge under prolonged or high-frequency demand rather than in the brief, submaximal efforts that behavioral testing requires. We employed gait analysis and beam walking as the measures best suited to early motor coordination, but these assess brief, voluntary movement and are therefore unlikely to reveal an impairment that depends on sustained activity. Endurance paradigms come closest to imposing prolonged demand, but are generally applied to adult mice, by which age glial coverage has largely been restored (Figure 4), so a deficit confined to earlier ages could escape detection. A decrement in force during repetitive nerve stimulation would test this prediction more directly than any behavioral measure. The absence of an overt behavioral phenotype therefore does not preclude a functional consequence of tSC loss, and may instead reflect the limits of what these measures can detect.

Our long-term analysis shows that glial coverage at the NMJ is eventually restored, as has been reported after tSC ablation using the Plp1-DTR model in adult muscle (Hastings et al., 2020). This suggests that the NMJ can sense the absence of glial coverage and trigger the migration and differentiation of neighboring aSCs to restore the niche. While gross motor function remained stable, body weight was reduced in tSC-ablated males but not females. We interpret this as a cumulative effect of the synaptic inefficiencies present at 4 weeks. Given that male mice undergo more rapid somatic growth, a homeostatic deficit confined to this period may impose a metabolic cost that is not subsequently recovered, and that manifests as reduced body weight.

In conclusion, our study establishes *Col20a1* as a genetic handle for tSC research. Rather than being required to hold the synapse together, tSCs appear to be required to keep it supplied: the mammalian NMJ structure is remarkably resilient to glial loss, while the efficiency of transmission during sustained activity depends on tSC-mediated regulation. With inducible genetic access largely restricted to these cells, this model provides a basis for dissecting tSC function and offers a framework for understanding glial-neuronal interactions and their contributions to neuromuscular health and disease.

## Materials and Methods

### Experimental Animals

The *Col20a1^CreERT2^* knock-in mice were generated on a C57BL/6N genetic background at the Lee Gil Ya Cancer and Diabetes Institute Center of Animal Care and Use, Gachon University (Incheon, Republic of Korea). The wild type C57BL/6J (RRID:ISMR_JAX:000664) mice, the *Rosa26^LSL-tdTomato^* (*Rosa*-tdT) (RRID:ISMR_JAX:007914) and the *Rosa26^LSL-DTA^* (*Rosa*-DTA) (RRID:ISMR_JAX:009669) reporter lines were acquired from the Jackson Laboratory (Bar Harbor, ME, USA) and maintained on a C57BL/6J genetic background. To generate *Col20a1^CreERT2^*; *Rosa26^LSL-tdTomato^* (*Col20a1*-tdT) mice, *Col20a1^CreERT2/+^* mice were crossed with *Rosa26^LSL-tdTomato/+^*, and both *Col20a1^+/+^*; *Rosa26^LSL-tdTomato/+^* and *Col20a1^CreERT/+^*; *Rosa26^LSL-tdTomato/+^* mice were used in experiments. To generate *Col20a1^CreERT2^*; *Rosa26^LSL-tdTomato/LSL-DTA^* mice (*Col20a1*-tdT/DTA), *Col20a1^CreERT2/+^*; *Rosa26^LSL-tdTomato/+^* mice were crossed with *Rosa26^LSL-DTA/+^*mice, generating both *_Col20a1CreERT2/+_*_; *Rosa26*_*_LSL-tdTomato/+_* _mice and *Col20a1*_*_CreERT2/+_*_; *Rosa26*_*_LSL-tdTomato/LSL-DTA_* _mice as_ littermates. In experiments using *Col20a1^CreERT2/+^*; *Rosa26^LSL-tdTomato/LSL-DTA^*mice, *Col20a1^CreERT2/+^*; *Rosa26^LSL-tdTomato/+^* mice were used as controls. All animal experiments were conducted in accordance with the ethical guidelines and approved by the Institutional Animal Care and Use Committee (IACUC) of Seoul National University (Approval No. SNU-250324-2-1). Mice were housed in a specific pathogen-free (SPF) facility under a 12-hour light/dark cycle with *ad libitum* access to standard chow and water. Both male and female mice were used in this study.

### Generation of *Col20a1*-CreERT2 Knock-in Mice

To generate the terminal Schwann cell (tSC)-specific *Col20a1*-CreERT2 knock-in mouse line, we employed a microinjection-free, CRISPR/Cas9-mediated homology-directed repair (HDR) strategy termed CRISPR-READI (CRISPR RNP electroporation and AAV donor infection)(Chen et al., 2019)Chen et al., 2019). A targeting donor vector was designed to insert a cassette containing a P2A self-cleaving peptide, the CreERT2 recombinase sequence, and an SV40 polyadenylation (pA) signal immediately upstream of the endogenous STOP codon in exon 35 of the *Col20a1* gene. To achieve high-efficiency integration, the HDR donor sequence was flanked by AAV2 inverted terminal repeats (ITRs) and packaged into recombinant AAV virions. Briefly, fertilized mouse zygotes were harvested and infected with the recombinant AAV harboring the HDR donor template. Subsequently, pre-assembled Cas9/single-guide RNA (sgRNA) ribonucleoproteins (RNPs) were introduced into the AAV-transduced zygotes via electroporation to induce DNA double-strand breaks and stimulate homologous recombination. Following successful homologous recombination, the targeted allele (*Col20a1*^CreERT2^) allows the co-translation of the endogenous COL20A1 protein and the CreERT2 recombinase, separated by the P2A peptide. Founder mice were identified by PCR genotyping and subsequently backcrossed with C57BL/6J mice to establish a stable germline-transmitted colony.

### Tamoxifen Administration for Inducible Labeling and Ablation

To induce Cre-mediated recombination, tamoxifen (Sigma-Aldrich) was dissolved in corn oil at a concentration of 20 mg/mL. For adult mice (8–12 weeks old), tamoxifen was administered via oral gavage at a dose of 160 mg/kg body weight for five consecutive days. For postnatal induction (e.g., P10), tamoxifen was administered via a single intraperitoneal (IP) injection at the same weight-adjusted dose (160 mg/kg). All mice including control mice lacking the CreERT2 allele received the identical tamoxifen regimen.

### Genotyping

Routine genotyping of the transgenic and mutant mice was performed by PCR using genomic DNA extracted from tail biopsies or ear snips. The tissues were lysed in DirectPCR Lysis Reagent (Viagen Biotech) supplemented with Proteinase K at a final concentration of 0.2 µg/µL. The lysis mixture was incubated at 65°C for 30 minutes, followed by a heat-inactivation step at 95°C for 1 hour.

The following primers were used: Col20a1_common_F: 5’ – ATC AGC CAC ACA AGC AAC CC – 3’, Col20a1_WT_R: 5’ – ATT GTT CAG CAG AGC TCC CC – 3’, Col20a1_Tg_R: 5’ – ACC GGC AAA CGG ACA GAA GC – 3’, Rosa-tdT_WT_F: 5’ – AAG GGA GCT GCA GTG GAG TA – 3’, Rosa-tdT_WT_R: 5’ – CCG AAA ATC TGT GGG AAG TC – 3’, Rosa-tdT_Tg_F: 5’ – GGC ATT AAA GCA GCG TAT CC – 3’, Rosa-tdT_Tg_R: 5’ – CTG TTC CTG TAC GGC ATG G – 3’, DTA_F: 5’ – GAC TGA CGA AGG TTC TCG CA – 3’, DTA_R: 5’ – GCT TAA CGC TTT CGC CTG TT – 3’.

PCR amplification was carried out using EmeraldAmp GT PCR Master Mix (Takara Bio) in a SimpliAmp Thermal Cycler (Applied Biosystems). For the *Col20a1*and *Rosa26*-tdTomato loci, the respective WT and Tg primer sets were combined in single multiplex PCR reactions. The general thermal cycling conditions are as follows: an initial denaturation at 95°C for 5 minutes, followed by 33 cycles of denaturation at 95°C for 15 seconds, annealing for 20 seconds, and extension at 72°C for 30 seconds, with a final extension at 72°C for 7 minutes. The annealing temperatures were individually optimized for each target: 65°C for the *Col20a1*-CreERT2 multiplex, 61°C for the *Rosa26*-tdTomato multiplex, and 57°C for the DTA amplification. Finally, the amplicons were resolved on 1.5% agarose gels, stained with EcoDye (Biofact), and visualized using a Davinch Gel Imaging System (Davinch-K).

### Single-cell RNA Sequencing Data Analysis

#### Data Acquisition and Quality Control

To characterize the transcriptomic profile of tSCs and other skeletal muscle-resident populations, we utilized a publicly available single-cell RNA sequencing (scRNA-seq) dataset generated from adult wild type mouse hindlimb muscles (Giordani et al., 2019), accessible via the NCBI Gene Expression Omnibus (GEO) under accession number GSE110878. All computational and statistical analyses were performed using R version 4.5.3 and the Seurat package version 5.4.0 (Hao et al., 2024). The datasets of two mice were read and further processed independently. Initial quality control was performed to filter out cells with fewer than 500 or more than 4,000 expressed genes, fewer than 500 or more than 9,000 total UMI counts, or greater than 10% mitochondrial gene expression.

#### Doublet Detection and Filtering

To facilitate doublet detection, standard preprocessing—including normalization, scaling, principal component analysis (PCA), and clustering—was initially performed on each sample independently. Specifically, the gene expression matrices were normalized using global-scaling normalization (“LogNormalize”) with a scale factor of 10,000. To identify highly variable features, the “vst” method was applied to select the top 2,000 highly variable genes. The data were then linearly scaled and centered. PCA was performed on the scaled data using the identified highly variable genes. To cluster the cells, a K-nearest neighbor (KNN) graph was constructed based on the Euclidean distance in PCA space, using the first 30 principal components (PCs). Cell clustering was performed using the Louvain algorithm with a resolution of 0.5. The DoubletFinder package was then applied to each dataset. The expected doublet rate was set to 4.9%. The optimal pK values were empirically determined as 0.06 and 0.005 for Sample 1 and Sample 2, respectively, utilizing a pN value of 0.25 and the first 30 PCs. The proportion of homotypic doublets was estimated based on the initial clustering to adjust the final doublet expectations. Only cells classified as “Singlet” were retained for subsequent analyses.

#### Data Integration and Dimensionality Reduction

The singlet-filtered objects from both samples were merged into a single dataset. To normalize the data, stabilize variance, and mitigate the confounding effects of mitochondrial gene expression, we applied the SCTransform method, explicitly regressing out the mitochondrial percentage (percent.mt). PCA was performed on the SCTransform-normalized data. Cell clustering was conducted using a KNN graph constructed on the first 30 PCs and the Louvain algorithm with a resolution of 0.5. Uniform Manifold Approximation and Projection (UMAP) was generated using the same top 30 PCs for non-linear dimensional reduction and visualization.

#### Schwann Cell Subsetting and Sub-clustering

The pan-Schwann cell population was initially annotated based on the enrichment of previously established marker genes, including *Plp1* and *S100b*, and subsequently subsetted for downstream analysis. The subsetted Seurat object was re-processed by normalizing with global-scaling (“LogNormalize”), scaling, and re-identifying variable features. PCA was re-calculated on the top 2,000 highly variable genes of this subpopulation. Sub-clustering was performed using the first 10 PCs and a KNN graph, followed by the Louvain algorithm with a resolution of 1.0. UMAP was generated using the same 10 PCs for visualization, which yielded distinct Schwann cell sub-clusters. The tSC population was annotated based on the enrichment of previously established marker genes (Castro et al., 2020), explicitly separating them from myelinating Schwann cells (mSCs) and Remak non-myelinating Schwann cells (nmSCs). Expression of selected genes was visualized using the FeaturePlot, VlnPlot and DotPlot functions in Seurat.

#### Differential Expression and Marker Gene Identification

To identify highly specific marker genes for putative tSCs, differential expression analysis was conducted using the FindMarkers function in Seurat, utilizing the non-parametric Wilcoxon rank-sum test. A two-tier intersection strategy was employed to ensure the identification of both highly representative and exclusively specific markers. First, differentially expressed genes (DEGs) were calculated by comparing the putative tSC cluster with the other Schwann cell sub-clusters. Significant DEGs for this lineage-specific comparison were filtered based on a log2 fold change > 1, an adjusted *p*-value < 0.05, and min.pct = 0.5. Secondly, a subsequent comparison was conducted between the putative tSC cluster and all the other cell populations in the whole-muscle dataset. For this tissue-wide comparison, alongside the same fold change and significance thresholds, significant DEGs were strictly filtered based on minimal expression in background cells (pct.2 < 0.01). The final tSC marker gene candidates were established by intersecting the significant DEGs derived from these two independent criteria.

#### Tissue-wide Expression Profiling

To assess the tissue-wide expression profiles and cellular specificity of the candidate genes across diverse organ systems, we utilized the Chan Zuckerberg CELLxGENE Discover platform (Program et al., 2025). Single-cell transcriptomic datasets spanning various adult mouse tissues and organs were acquired to evaluate the expression patterns of the selected candidate genes, including *Col20a1*, *Acsbg1*, and *Fabp7*. Cell-type-specific enrichment patterns were analyzed and visualized as dot plots in R.

### Analysis of RNA-seq Datasets

To cross-validate and robustly identify tSC-specific marker genes, we performed a meta-analysis utilizing publicly available RNA-sequencing (RNA-seq) datasets from previous studies. Transcriptomic data profiling isolated synaptic Schwann cells (Castro et al., 2020) and skeletal muscle/NMJ tissues (Ham et al., 2020) were retrieved from the NCBI Gene Expression Omnibus (GEO) database. The datasets are accessible under the GEO accession numbers GSE152774 and GSE139209, respectively. Raw count matrices were downloaded for downstream computational analysis. The normalized read counts, expressed as Counts Per Million (CPM) or Transcripts Per Million (TPM), for each candidate gene and for *Kcnj10* (Figure 1—figure supplement 5) were visualized using GraphPad Prism software (version 9, GraphPad Software). The DEG analysis for bulk RNA-seq data were performed within R using DESeq2 package version 1.50.2. For the DESeq2 workflow, raw counts were normalized using the median of ratios method to account for sequencing depth variations, and gene-wise dispersion estimates were modeled based on a negative binomial distribution. To identify DEGs, the Wald test was applied, and *p*-values were adjusted for multiple testing using the Benjamini-Hochberg false discovery rate (FDR) procedure. Genes were defined as significantly differentially expressed if they met the strict criteria of an adjusted *p*-value (FDR) < 0.05 and a log2 fold change > 1. Volcano plots for DEG visualization were generated using ggplot2 packages in R.

### Isolation and Sorting of Muscle-Resident Cells

To isolate tdTomato labeled (tdT^+^) cells and categorize distinct populations of muscle-resident cells from tamoxifen-treated *Col20a1*-tdT mice, fluorescence-activated cell sorting (FACS) was performed based on a previously described protocol (Liu et al., 2015) with minor optimizations. Briefly, hindlimb muscles were harvested, finely minced using surgical scissors, and rinsed in Dulbecco’s Modified Eagle Medium (DMEM; HyClone) supplemented with 10% horse serum (Gibco). For enzymatic tissue digestion, the minced samples were incubated in DMEM containing 10% horse serum, 800 U/mL Collagenase II (Worthington Biochemical), and 1.1 U/mL Dispase II (Gibco) for 40 minutes at 37°C with gentle agitation. Subsequent mechanical dissociation was achieved by triturating the suspension through a 20-gauge needle ten times. The resulting homogenate was passed through a 40-μm cell strainer to obtain a mononuclear single-cell suspension. For immunophenotyping, the cells were incubated with the following primary antibodies (all diluted at 1:100 in PBS containing 10% horse serum): APC-conjugated anti-mouse CD31, CD45, and Ter119 (Biolegend); biotin-conjugated anti-mouse VCAM1 (BioLegend); and FITC-conjugated anti-mouse Sca-1 (BD Pharmingen). To detect the biotinylated VCAM1 antibody, cells were secondarily stained with PE/Cy7-conjugated streptavidin (1:100 in PBS containing 10% horse serum). Cell viability was assessed by adding 7-aminoactinomycin D (7-AAD; Sigma-Aldrich) to exclude dead cells. Following the staining process, the cells were resuspended in a FACS buffer consisting of 2% horse serum in PBS immediately prior to analysis.

Cell sorting was executed using a BD FACS Aria III cell sorter (BD Biosciences) controlled by BD FACS Diva software for data acquisition. Live (7-AAD^-^) cells were gated and sorted into specific sub-populations using the following strategies: muscle stem cells (MuSCs; Lin^−^/VCAM1^+^/Sca-1^−^), fibro-adipogenic progenitors (FAPs; Lin^−^/VCAM1^−^/Sca-1^+^), linage-positive cells (Lin^+^; CD31^+^, CD45^+^ or Ter119^+^), and double-negative cells (DN; Lin^−^/VCAM1^−^/Sca-1^−^). Within each gated population, tdT^+^ cells were quantified. Finally, tdT^+^ and tdT^−^ fractions from the DN population were independently sorted to evaluate genomic DNA recombination via PCR. Post-sorting data analysis and structural visualization were performed using FlowJo software (Tree Star).

### Genomic DNA Extraction and Recombination Analysis

To verify the genomic recombination of the *Rosa26*-LSL-tdTomato allele in specific muscle-resident cell populations, FACS-sorted cells were subjected to genomic DNA (gDNA) extraction using a standard phenol-chloroform method. Briefly, the sorted cells were lysed and homogenized in 100 µL of Tris-EDTA (TE) buffer containing 1% SDS, followed by overnight incubation at 50°C. For complete protein digestion, Proteinase K was added to a final concentration of 0.1 µg/µL, and the lysates were incubated at 37°C for 1 hour. The samples were adjusted to a volume of 250 µL with TE buffer and extracted with an equal volume of phenol-chloroform-isoamyl alcohol (25:24:1, Invitrogen). Following centrifugation at 20,000 × g for 3 minutes at 4°C, the upper aqueous phase was carefully collected. The gDNA was precipitated by adding 0.1 volumes of sodium acetate (2.5 M, pH 5.2) and 2 volumes of absolute ethanol, followed by incubation at −80°C for at least 1 hour. The DNA pellets were recovered by centrifugation (20,000 × g for 15 minutes at 4°C), washed twice with 70% ethanol, completely air-dried, and resuspended in TE buffer at 60°C.

To detect the tdTomato recombination event, PCR was performed using the purified gDNA as a template. The specific primer sequences used for the amplification were as follows: CAG_tdTomato_F, 5’-GCAACGTGCTGGTTATTGTG-3’ and CAG_tdTomato_R, 5’- CGCATGAACTCTTTGATGACC-3’. The PCR amplification was carried out using EmeraldAmp GT PCR Master Mix (Takara Bio) in a SimpliAmp Thermal Cycler (Applied Biosystems). The thermal cycling conditions consisted of an initial denaturation at 95°C for 5 minutes, followed by 33 cycles of denaturation at 95°C for 15 seconds, annealing at 56°C for 20 seconds, and extension at 72°C for 30 seconds. This was followed by a final extension step at 72°C for 7 minutes. The expected size of the recombined amplicon was exactly 280 bp. Finally, the PCR products were electrophoretically separated on a 1.5% agarose gel, stained with EcoDye nucleic acid staining solution (Biofact), and visualized using a Davinch Gel Imaging System (Davinch-K).

### Whole-Mount Immunofluorescence

#### Tissue Preparation and Whole-Mount Staining

For the majority of histological analysis, the extensor digitorum longus (EDL) muscle was primarily used. For the comparative analysis across diverse muscle types (Figure 2C), additional skeletal muscles (masseter, sternomastoid, diaphragm, psoas, and soleus) were also harvested and processed in the same manner. All freshly isolated muscles were immediately fixed in 4% paraformaldehyde (PFA) in PBS for 15 minutes at room temperature. Fixed muscles were quenched in 0.1M glycine in PBS for 1 hour at room temperature. After rinsing with PBS, muscle bundles were teased into smaller fibers using fine forceps under a dissecting microscope. The teased muscle fibers were incubated in a blocking buffer containing 5% bovine serum albumin and 2% Triton X-100 in PBS for 2 hours at room temperature. Tissues were then incubated with primary antibodies diluted in the blocking buffer for 3 days at 4°C. The primary antibodies used in this study include: rabbit anti-S100 (Ready-to-use formulation further diluted at 1:5; Agilent, IR50461-2) and mouse anti-neurofilament (1:1000; DSHB, 2H3). After washing three times with washing buffer (2% Triton X-100 in PBS) for 1 hour each, the tissues were incubated with a mixture of secondary antibodies and specific fluorescent probes diluted in blocking buffer for 3 days at 4°C. To visualize targeted proteins and postsynaptic acetylcholine receptors (AChRs), the following secondary antibodies and probes were used as dictated by the experimental design: Alexa Fluor 647-conjugated goat anti-rabbit IgG (1:500; Invitrogen, A32733), Alexa Fluor 405-conjugated goat anti-mouse IgG (1:500; Invitrogen, A31553), Alexa Fluor 488-conjugated α-bungarotoxin (α-Btx, 1:500; Invitrogen, B13422), and/or Hoechst 33342 (1:2000; Thermo Fisher, 62249). Following three 1-hour washes in washing buffer, the stained muscle fibers were whole-mounted on glass slides using Vectashield without DAPI (Vector Laboratories, H-1000).

#### Confocal Image Acquisition

Fluorescence images of the whole-mount stained muscles were acquired using a Leica TCS SP8 confocal laser scanning microscope equipped with 40× and 63× water-immersion objective lenses. Z-stack images were collected encompassing the entire depth of the motor endplate band. The laser power, gain, and pinhole settings were kept consistent across all biological replicates to ensure comparability. Image processing was performed using LAS X software (Leica Microsystems). Z-stack confocal images were collapsed into maximum intensity projections (MIPs). The structural integrity of the NMJs, including the degree of postsynaptic AChR cluster fragmentation and the status of motor axon innervation, was assessed via visual inspection of the MIPs. Multiple fields of view from at least three independent biological replicates were evaluated to determine the predominant morphological phenotypes.

#### Classification of Glial Coverage at NMJs

For the time-course analysis of tSC ablation and recovery (Figure 4), EDL muscles were collected at P14, P17, P21 and at 4, 6, 8 and 12 weeks of age, with three mice per time point. All NMJs within the imaged endplate band were evaluated, with no selection applied, by an investigator blinded to genotype and time point, using images acquired and displayed with identical settings. The tdTomato and S100B signals were assessed independently for every α-Btx-labeled NMJ, so that glial coverage could be judged without reference to the reporter. Each NMJ was assigned to one of four categories according to the S100B signal within the α-Btx-labeled area: reporter-positive NMJs carrying S100B-positive tSCs (tdT^+^ S100B^+^); and reporter-negative NMJs with no S100B signal (tdT^−^ S100B absent), with S100B leaving part of the area uncovered (tdT^−^ S100B partial), or with S100B covering it entirely, as in controls (tdT^−^ S100B full). Category proportions were calculated per animal and used as the unit of analysis; 3,087 NMJs were scored in total.

### Electrophysiology

The extensor digitorum longus (EDL) muscles, together with the attached peroneal nerve, were carefully dissected from mice and pinned to a Sylgard-coated recording chamber. Intracellular recordings were performed using 3 M KCl-filled glass microelectrodes submerged in oxygenated Ringer’s solution (138.8 mM NaCl, 4 mM KCl, 12 mM NaHCO_3_, 1 mM KH_2_PO_4_, 1 mM MgCl_2_ and 2 mM CaCl_2_ with a pH of 7.4). During the initial dissection process, Ca^2+^ free Ringer’s solution was utilized to prevent muscle contraction and maintain structural stability. The evoked endplate potentials (eEPPs) were elicited using bipolar stimulation of the peroneal nerve. Prior to recording, muscle action potentials were completely blocked by incubating the preparation with 2.5 μM μ-conotoxin GIIIB (Alomone, Jerusalem, Israel) for 10 minutes. Synaptic transmission efficacy was evaluated using paired-pulse stimulation (with 10, 20, and 50 ms inter-stimulus intervals) and high-frequency train stimulation (20 pulses at 20 Hz). Data were acquired and analyzed using an Axoclamp 900A amplifier, a Digidata 1550B digitizer, and Clampfit 10.7 software.

For compound muscle action potential (CMAP) measurement, mice were fully anesthetized via an intraperitoneal injection of Avertin (250 mg/kg; Sigma-Aldrich). The sciatic nerve was percutaneously stimulated at the sciatic notch using an Isolated Pulse Stimulator Model 2100 (A-M Systems). Supramaximal square-wave stimuli (∼70 mA) were delivered at a frequency of 1 Hz with a pulse duration of 0.1 ms. For signal acquisition, a recording electrode was carefully inserted subdermally over the belly of the gastrocnemius (GA) muscle, ensuring that the underlying muscle fibers were not physically punctured. Reference and ground electrodes were positioned near the Achilles tendon and on the tail, respectively. Electromyographic signals were captured using an IX-RA-834 Data Recorder (iWorx). The CMAP amplitude was quantified by calculating the absolute voltage difference between the maximum positive and negative peaks. For robust statistical analysis, four consecutive CMAP traces were elicited from each GA muscle, and the mean amplitude of these measurements was utilized as the representative value for each animal.

### Electron Microscopy

#### Tissue Processing and OTO Staining

EDL muscles were dissected from mice and immediately fixed in 2.5% glutaraldehyde and 2% paraformaldehyde in 0.1 M sodium cacodylate buffer overnight at 4°C. To enhance the electron contrast of lipid membranes and synaptic vesicles, the tissues were processed using an OTO (osmium-thiocarbohydrazide-osmium) staining protocol. Samples were post-fixed in a solution of 1% osmium tetroxide and 1.5% potassium ferrocyanide for 1 hour at room temperature. Then, samples were incubated in a 1% thiocarbohydrazide (TCH) solution for 20 minutes, followed by incubation in a 2% osmium tetroxide solution for 30 minutes at room temperature. Following *en bloc* staining with 0.5% uranyl acetate overnight at 4°C, samples were further stained with 20 mM lead aspartate in a 60°C oven, and subsequently dehydrated through a graded ethanol series (30%, 50%, 70%, 80%, 90%, and 100% three times, 20 minutes each at room temperature). Following dehydration, samples were cleared in 100% propylene oxide (PO) for 15 minutes twice. For resin infiltration, the samples were incubated in a 1:1 mixture of PO and Spurr’s resin (Electron Microscopy Sciences) for 1.5 hours, followed by a 1:2 mixture of PO and Spurr’s resin for an additional 1.5 hours. Finally, the samples were immersed in 100% pure Spurr’s resin and incubated overnight at room temperature for complete infiltration. The 100% Spurr’s resin was changed the next day, and the samples were incubated for an additional 3 hours. Finally, the samples were embedded in molds with fresh resin and polymerized in an oven at 70°C.

#### Ultramicrotome Section and BSE-SEM Imaging

Embedded resin blocks were trimmed, and ultrathin sections with a thickness of 70 nm were cut using an ultramicrotome (Leica EM UC7). The ultrathin sections were carefully collected onto conductive silicon wafers. High-resolution imaging of the presynaptic terminals was performed using a SIGMA 360 Scanning Electron Microscope (SEM; Carl Zeiss). Images were acquired utilizing a backscattered electron detector (BSD). The microscope was operated at an accelerating voltage of 3.0 kV with a working distance of 6.5 mm.

#### Quantitative Morphometric Analysis of the NMJ

Quantitative morphometric analysis of the neuromuscular junctions (NMJs) was performed using ImageJ/Fiji software (NIH) by an investigator blinded to the genotypes. For each analyzed NMJ, the cross-sectional area of the presynaptic motor terminal was delineated by manually tracing the presynaptic membrane in the SEM-BSD images. All clearly distinguishable synaptic vesicles within this defined presynaptic profile were counted. The synaptic vesicle density was calculated by dividing the total number of counted vesicles by the cross-sectional area of the presynaptic terminal, expressed as vesicles per square micrometer (vesicles/µm²). The synaptic cleft width was determined by measuring the distance between the pre- and postsynaptic membranes at 7 to 10 distinct, regularly spaced points per NMJ and averaging the values. The number of junctional folds was counted and normalized to the synaptic apposition length. To examine the spatial distribution of synaptic vesicles, the perpendicular distance from the presynaptic membrane apposed to the postsynaptic membrane to the center of each vesicle was measured. Vesicles lying within 50 nm of this membrane were defined as the membrane-proximal pool, and their number was normalized to the length of the apposed presynaptic membrane (vesicles/µm). For the distribution analysis, vesicles were grouped by the same distance measure into ≤100, 100–200 and >200 nm bins, and the number of vesicles in each bin was normalized to the cross-sectional area of the presynaptic terminal (vesicles/µm²). In total, 23 NMJ profiles from 6 mice (control), 15 NMJ profiles from 6 mice (tSC-retained) and 23 NMJ profiles from 5 mice (tSC-ablated) were analyzed.

### Behavioral Assays

#### General Behavioral Testing Procedures

All behavioral assays were performed during the light cycle, at a fixed time of the day (3 PM). To minimize stress, mice were transferred to the behavioral testing room and allowed to acclimatize for at least 30 minutes prior to the onset of any experiments. To ensure objectivity, all behavioral tests and subsequent data analyses were conducted by an investigator strictly blinded to the genotypes of the mice. Both male and female mice were used in the experiments, and no sex difference was observed in any of the behavioral tests.

#### Gait Analysis

To evaluate the walking pattern and locomotor coordination of 4-week-old mice, gait analysis was performed. The forepaws and hindpaws of the mice were coated with non-toxic ink of different colors. The mice were then allowed to walk along a paper-lined runway (50 cm long, 6 cm wide) into a dark goal box. From the recorded footprints, the stride length and stride width (base of support) were measured to assess motor coordination.

#### Beam Walking Test

Prior to the actual testing, 4-week-old mice underwent a pre-training phase to familiarize themselves with the apparatus. Mice were trained to cross an elevated cylindrical wooden beam (50 cm long, 3 cm diameter) elevated 30 cm above a padded surface, leading to an enclosed safety box. After completing three consecutive training trials, the mice were transitioned to a narrower testing beam (50 cm long, 1 cm diameter) for further adaptation. The actual test was conducted on the following day, using the testing beam. Each mouse was subjected to three independent trials, and the latency to successfully cross the beam was recorded for each trial. The values from the three individual trials were averaged to determine the representative crossing latency for each animal.

#### Grip Strength Test

Whole-body muscle strength was evaluated using a digital grip strength meter (BIO-GS3; Bioseb). 12-week-old mice were placed on the apparatus and allowed to firmly grasp the metal grid with all four paws. The base of the tail was gently and steadily pulled backward parallel to the grid until the animal released its grip. The maximum peak force exerted was recorded in Newtons (N). Each testing session consisted of five consecutive pulls. The three highest peak force values out of the five trials were averaged to determine the score for that session. Each animal underwent a total of three independent testing sessions, separated by a resting interval of at least 30 minutes. The overall absolute grip strength for each mouse was calculated by averaging the scores from these three sessions. The absolute peak force values were normalized to the individual body weight of each mouse to account for potential body size differences among the cohorts.

#### Rotarod Test

Motor coordination, balance, and physical endurance were assessed using an accelerating rotarod apparatus (LE8500, Panlab/Harvard Apparatus). Following a brief habituation phase to the testing room, mice were carefully placed on the rotating rod. The apparatus was programmed to accelerate linearly from an initial speed of 4 rpm to a maximum speed of 40 rpm over a 5-minute period. The trial was terminated when the mouse fell off the rod, and the latency to fall (in seconds) was automatically recorded. Each mouse was subjected to three independent trials with a minimum resting interval of 15 minutes. The final representative latency to fall for each mouse was determined by averaging the values obtained from the three trials.

#### Treadmill Endurance Running

Exercise capacity and endurance limits were assessed using a motorized rodent treadmill (DJ2-242 Dual Treadmill; Daejong Instrument) equipped with a mild electrical shock grid to provide running motivation. Prior to the actual testing, mice were acclimated to the treadmill apparatus for three consecutive days. Each daily training session consisted of a 5-minute stationary exploration period (0 m/min), followed by running at 5 m/min, 10 m/min, and 15 m/min for 5 minutes at each speed. On the actual test day, the treadmill was set at a 0° incline. The initial belt speed was set to 10 m/min and was progressively increased by 2 m/min every 20 minutes until the animals reached exhaustion. Exhaustion was strictly defined as the point at which a mouse remained on the shock grid for more than 10 consecutive seconds without attempting to resume running. Both the total running time and the total distance covered were recorded for each animal.

### Statistical Analysis

Statistical analyses for the histological and behavioral assays were performed using GraphPad Prism software (version 9, GraphPad Software). For electrophysiological experiments, statistical analysis was performed using SPSS Statistics software (version 29.0.1.0; IBM). All quantitative data were visualized using GraphPad Prism version 9 and presented as the mean ± standard deviation (SD), except for the electrophysiological data, which were expressed as the mean ± standard error of the mean (SEM). Prior to statistical comparisons, the normality of the data distribution was assessed using the Shapiro-Wilk test, and the homogeneity of variances was evaluated using the F-test or Brown-Forsythe test. For comparisons between two independent experimental groups, a two-tailed unpaired Student’s t-test was utilized for normally distributed data with equal variances. If the assumption of equal variances was violated, Welch’s t-test was applied. If the data did not meet the assumption of normality, a nonparametric Mann-Whitney U test was applied. For comparisons involving three or more groups, a one-way analysis of variance (ANOVA) followed by Bonferroni’s post hoc test was performed. For comparisons involving two factors, a two-way ANOVA followed by Šídák’s multiple comparisons test was performed. For the analysis of train data involving repeated measurements, a repeated measures two-way ANOVA followed by Bonferroni’s multiple comparisons test was performed. The time-course data in Figure 4 characterize the progression of glial coverage loss and recovery rather than testing specific contrasts and were therefore not subjected to statistical comparison. In the figures, exact *p-*values are shown for comparisons that reached statistical significance, which was defined as a *p*-value of less than 0.05. *p*-values are reported to a maximum of four decimal places and two significant figures. Values below 0.0001 are reported as *p* < 0.0001. Where comparisons were made at multiple points across a stimulus train, the result of the overall test is reported in the figure and individual comparisons are indicated by asterisks, as denoted as follows: \**p* < 0.05, \*\**p* < 0.01 and \*\*\**p* < 0.001.

## Data Availability Statement

The publicly available transcriptomic datasets analyzed in this study can be accessed in the NCBI Gene Expression Omnibus (GEO) under accession numbers GSE110878, GSE152774, GSE139209, and the CZ CELLxGENE Discover platform. All other data generated or analyzed during this study (including behavioral, electrophysiological, and morphological data) are included in the manuscript and its supporting files. Source data files will be provided for all quantitative figures upon request or prior to publication.

## Acknowledgements

This work was supported by grants from the National Research Foundation of Korea (NRF) funded by the Korean government (MSIT) (Grant Nos. RS-2025-00556439, RS-2020-NR049538, and RS-2026-25506025). The generation of the *Col20a1*-CreERT2 mouse model was supported by the NRF grant funded by the MSIT (Grant No. RS-2024-00400118). During the preparation of this work, the authors used Google Gemini (Gemini 1.5 Pro) in order to refine the English language and improve sentence readability. After using this tool/service, the authors reviewed and edited the content as needed and take full responsibility for the content of the publication.

## Author Contributions

H.K., K.Y., and Y.-Y.K. designed research; K.Y. discovered the marker gene; H.K. performed *in vivo* experiments, morphological, and behavioral research; S.-Y.K. performed electrophysiological research; S.-Y.C. contributed new reagents/analytic tools; H.K., S.-Y.K., K.Y., S.-Y.C., and Y.-Y.K. analyzed data; and H.K. and Y.-Y.K. wrote the paper.

---

**Figure 1—figure supplement 1.**
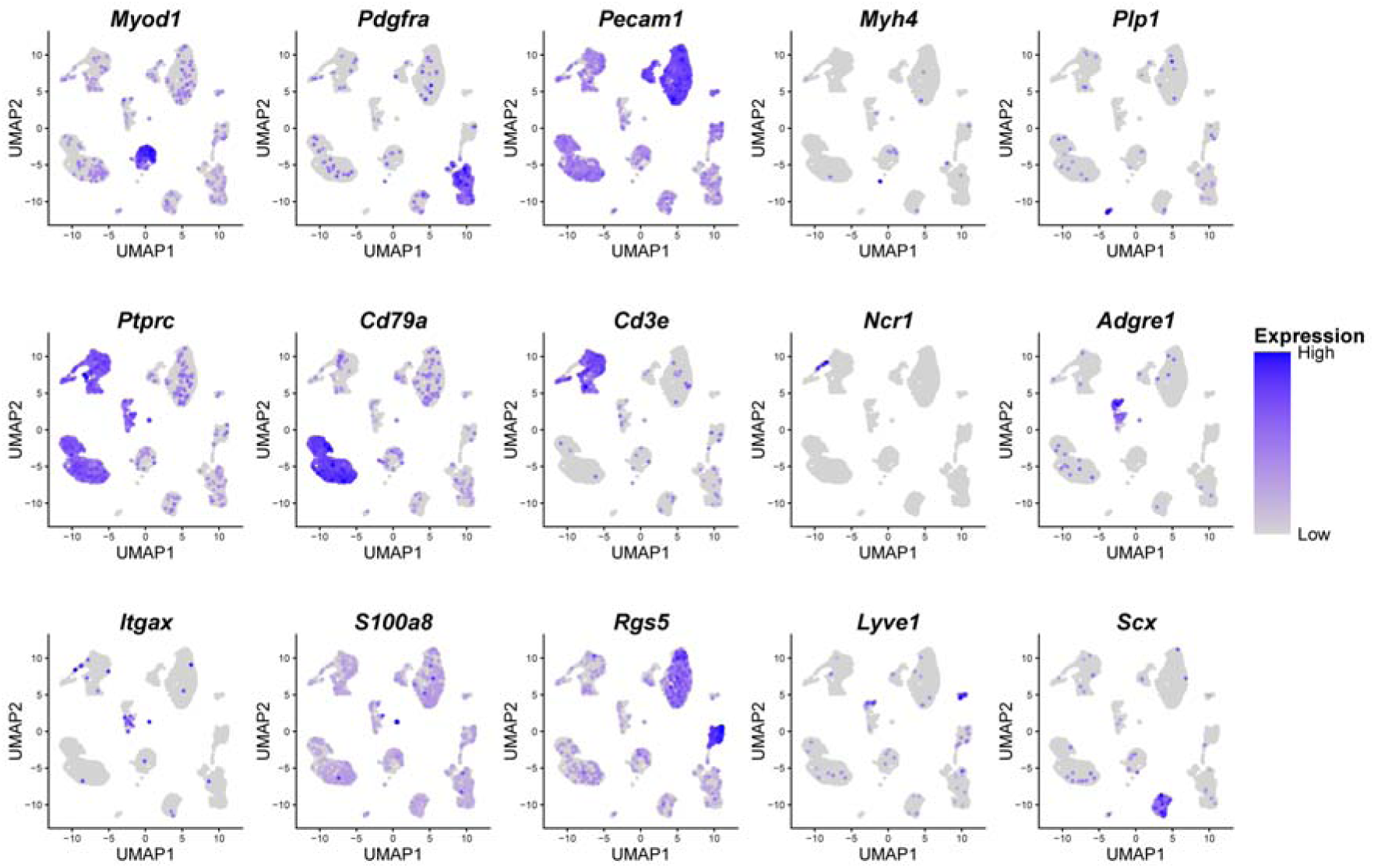
Expression profiling of canonical cell-type markers in the skeletal muscle-resident cell population. UMAP feature plots showing the distribution and expression levels of established marker genes for diverse cell types within the whole skeletal muscle scRNA-seq dataset.

**Figure 1—figure supplement 2.**
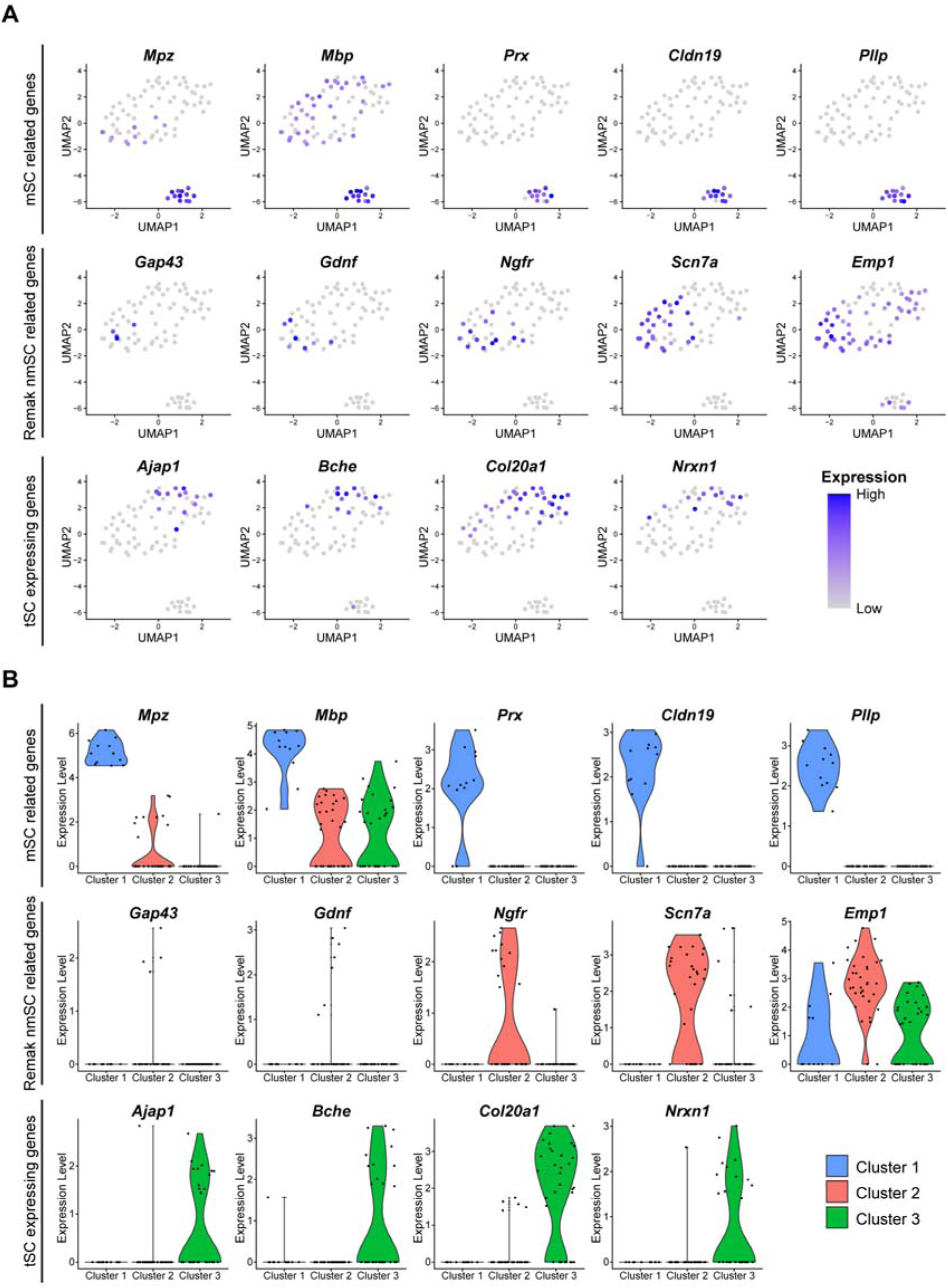
Validation of Schwann cell sub-cluster identities using established subtype-specific marker genes. (A) UMAP feature plots and (B) Violin plots illustrating the distinct expression patterns of known Schwann cell subtype markers across the three isolated sub-clusters.

**Figure 1—figure supplement 3.**
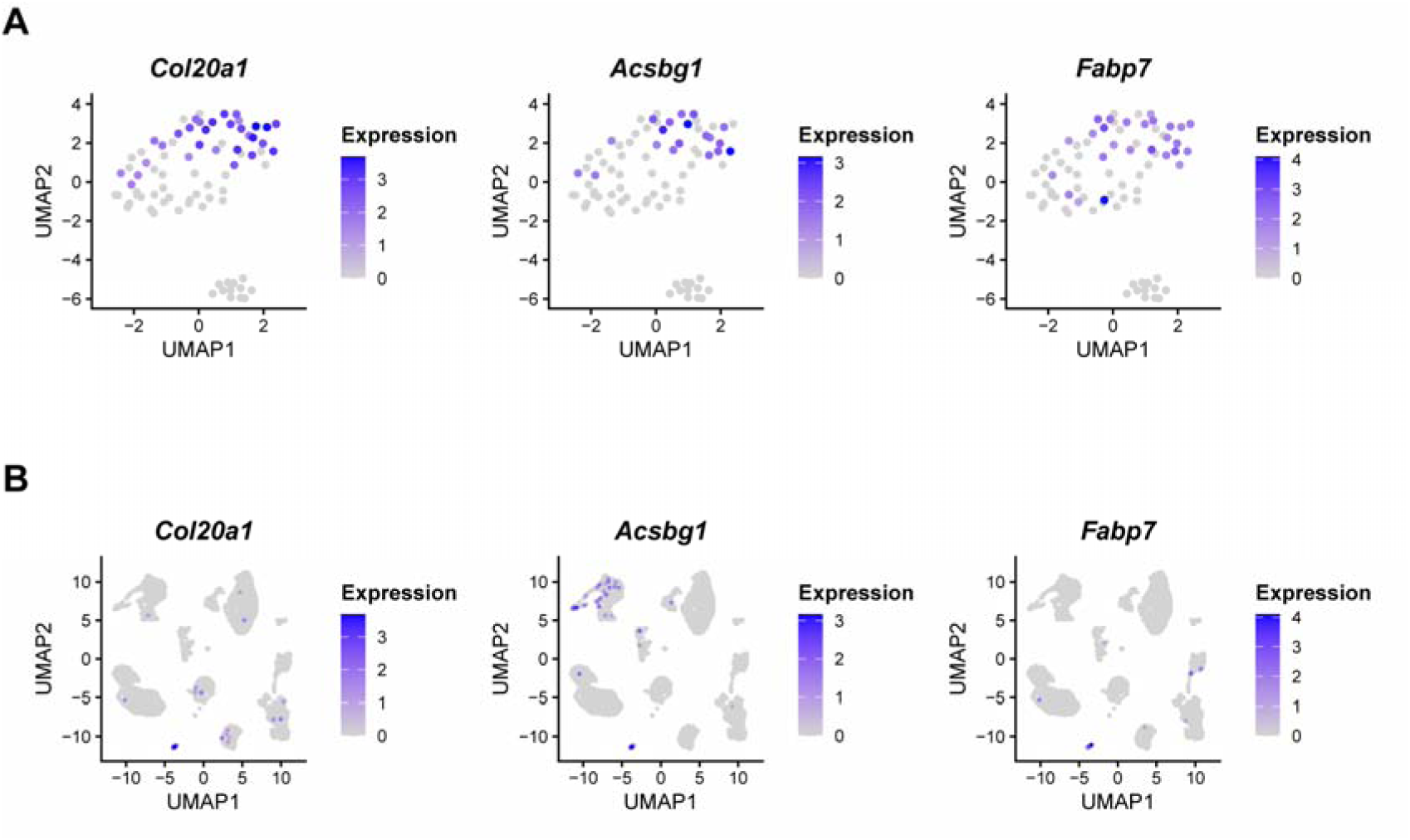
Highly restricted expression of the three final candidate genes within the tSC cluster. UMAP feature plots illustrating the localized expression patterns of the three selected candidate genes (*Col20a1*, *Acsbg1*, and *Fabp7*). The plots highlight their highly specific enrichment in the tSC population when mapped across (A) the isolated Schwann cell sub-clusters and (B) the entire skeletal muscle-resident cell population.

**Figure 1—figure supplement 4.**
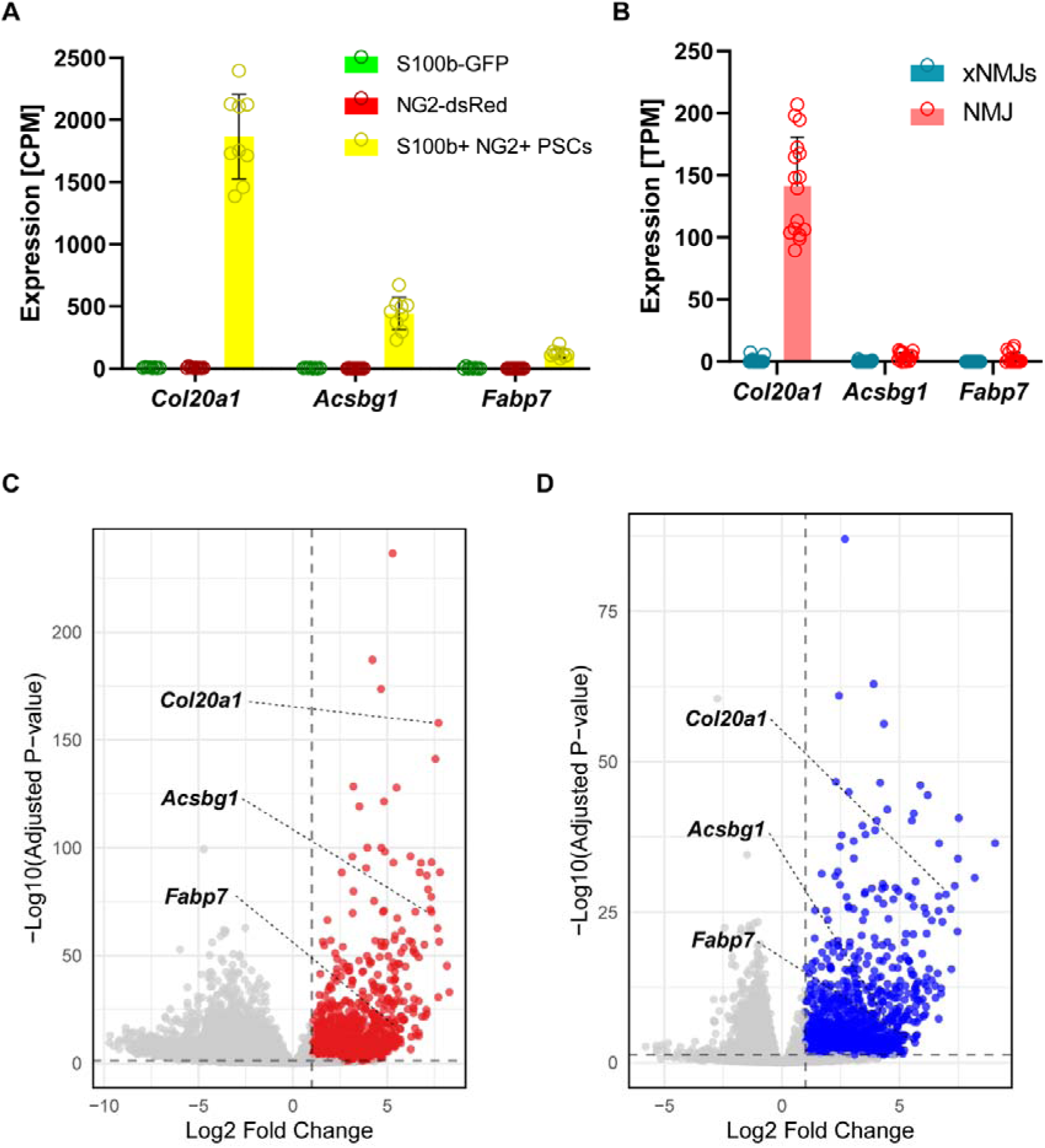
Independent bulk RNA-seq analyses confirm *Col20a1* as the most robust tSC-specific marker. Re-analysis of publicly available bulk RNA-seq datasets validates the enrichment of the candidate genes in tSCs. (A) Bar graphs showing expression levels (CPM) of the three candidate genes, *Col20a1*, *Acsbg1*, and *Fabp7*, in FACS-sorted S100b-GFP/NG2-dsRed double-positive tSCs compared to S100b-GFP or NG2-dsRed single positive cells (Castro et al., 2020). (B) Bar graphs showing expression levels (TPM) of the three candidate genes in micro-dissected NMJ regions (NMJ) compared to non-NMJ regions (xNMJ) (Ham et al., 2020). (C, D) Volcano plots indicating the statistical significance (–Log10 adjusted *p*-value) and Log_2_FC of the candidate genes derived from the datasets in (A) and (B), respectively. For dataset (A), the volcano plot in (C) displays the differential expression between double-positive tSCs and the combined single-positive cell populations. *Col20a1* exhibits the highest expression level and the most significant fold-change among the candidates, prioritizing it for the genetic model generation. Data in (A) and (B) are presented as mean ± SD. (A) n = 9 (S100b+/NG2+ PSCs), 8 (S100b-GFP), and 10 (NG2-dsRed) samples; (B) n = 16 samples per group. (A) and (B) display expression levels; the corresponding differential statistics are shown in (C) and (D), where significance was assessed with DESeq2 using the Wald test with Benjamini-Hochberg correction.

**Figure 1—figure supplement 5.**
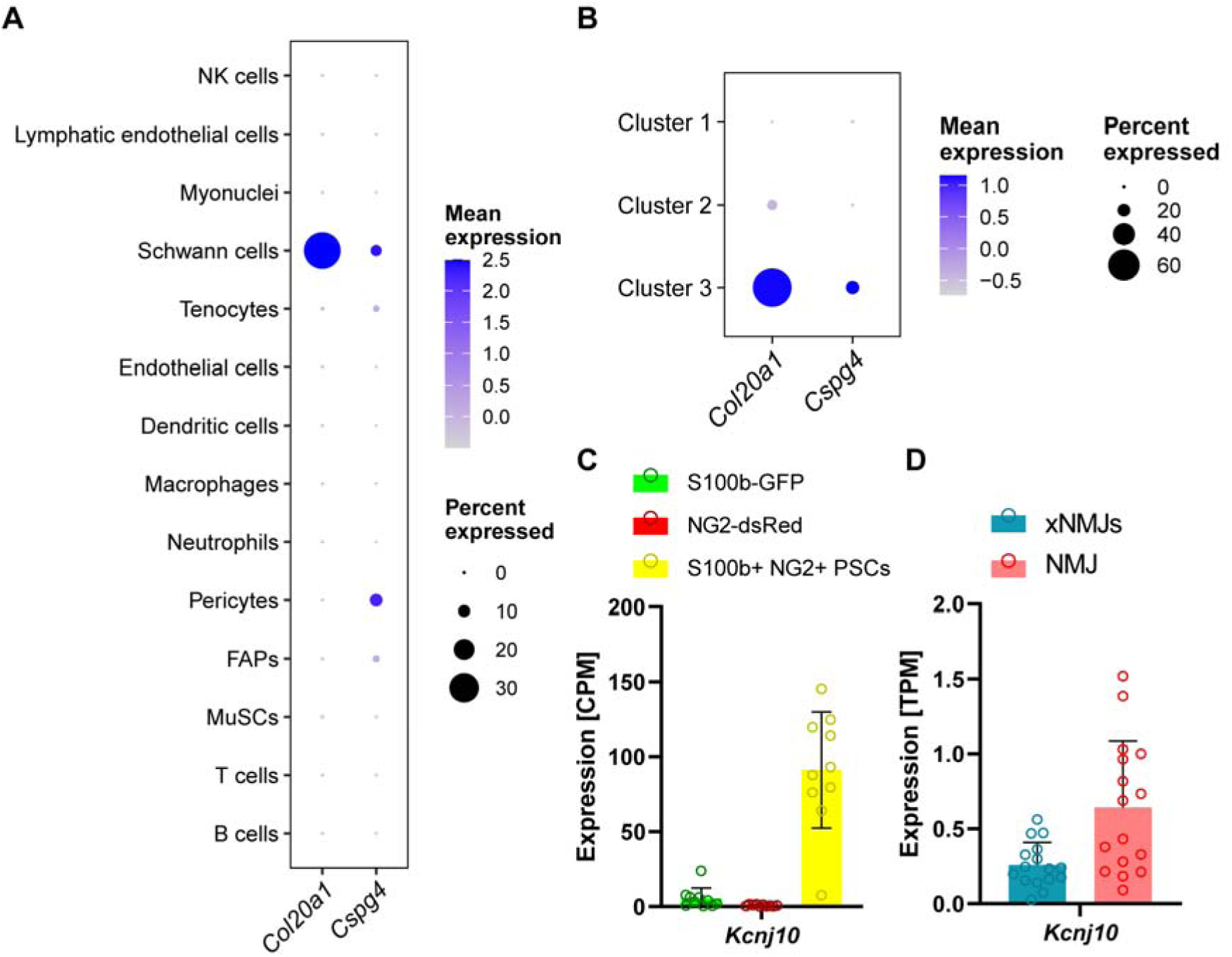
Expression of *Col20a1* relative to the established tSC markers *Cspg4* and *Kcnj10*. Comparison of *Col20a1* expression with that of *Cspg4* (NG2) and *Kcnj10* (Kir4.1), two genes previously reported to be expressed at tSCs (Castro et al., 2020; Procacci et al., 2023), using single-cell dataset analyzed in Figure 1 and publicly available bulk RNA-seq datasets re-analyzed in Figure 1—figure supplement 4. *Kcnj10* was not detected in the single-cell dataset and is therefore shown only for the bulk datasets. (A) Dot plot showing the expression of *Col20a1* and *Cspg4* (NG2) across the cell types of skeletal muscle. Node size indicates the percentage of cells expressing each gene and color intensity represents the mean expression level. *Col20a1* and *Cspg4* are both detected in Schwann cells, with *Cspg4* also detected in pericytes. (B) Dot plot showing the expression of the same genes across the three Schwann cell sub-clusters defined in Figure 1C. Within the Schwann cell compartment, both genes are enriched in Cluster 3, the cluster identified as tSCs in Figure 1. Color scales in A and B are set independently. (C) Expression of *Kcnj10* (CPM) in FACS-sorted S100b-GFP/NG2-dsRed double-positive tSCs compared with S100b-GFP or NG2-dsRed single-positive cells (Castro et al., 2020). *Kcnj10* is enriched in the double-positive population. (D) Expression of *Kcnj10* (TPM) in micro-dissected NMJ regions (NMJ) compared with non-NMJ regions (xNMJ) (Ham et al., 2020). *Kcnj10* is likewise enriched at NMJ regions, although its absolute expression level is low. *Col20a1* expression in the same bulk datasets is shown in Figure 1—figure supplement 4. Data in (C) and (D) are presented as mean ± SD. (C) n = 9 (S100b+/NG2+ PSCs), 8 (S100b-GFP), and 10 (NG2-dsRed) samples; (D) n = 16 samples per group. These panels display expression levels only, and no statistical comparison was performed between groups.

**Figure 2—figure supplement 1.**
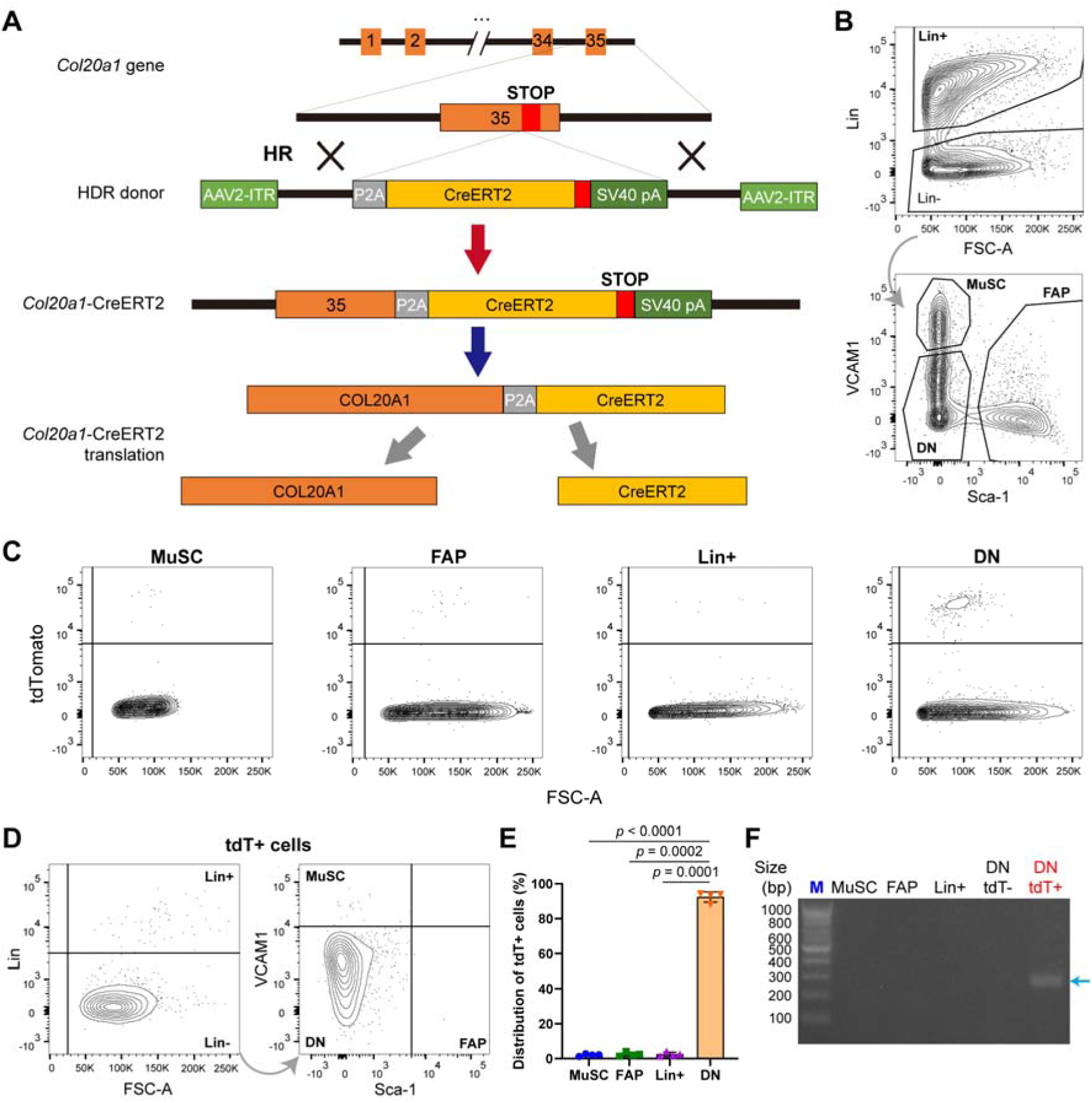
Generation strategy and quantitative flow cytometry validation of the *Col20a1*-CreERT2 mouse model. (A) Schematic diagram illustrating the design of the *Col20a1*-CreERT2 knock-in allele. A P2A-CreERT2 cassette was inserted at the C-terminus of the endogenous *Col20a1* gene via homologous recombination (HR). (B) Flow cytometry gating strategy for isolating mononuclear cells from whole skeletal muscle. Cells were separated into Lineage-positive (Lin^+^; CD31^+^, CD45^+^ or Ter119^+^), muscle stem cells (MuSC; Lin ^−^/VCAM1^+^/Sca-1^−^), fibro-adipogenic progenitors (FAP; Lin^−^/VCAM1^−^/Sca-1^+^), and double-negative cells (DN; Lin^−^/VCAM1^−^/Sca-1^−^), the latter of which harbors the Schwann cell population. (C) Representative flow cytometry plots displaying tdTomato fluorescence across the four isolated cell populations. (D, E) Flow cytometry analysis (D) and quantification (E) of the tdTomato-positive (tdT^+^) cell distribution. The vast majority of tdT^+^ cells (∼92%) are highly restricted to the DN population. Data are presented as mean ± SD (n = 4). Statistical significance was determined using Repeated Measures ANOVA with Bonferroni’s post hoc test. Exact *p*-values are shown for comparisons that reached statistical significance (*p* < 0.05). (F) PCR validation of genomic DNA extracted from the sorted cell populations. The recombined tdTomato amplicon (blue arrow) is exclusively detected in the tdT^+^ fraction of the DN population. M: DNA size marker.

**Figure 4—figure supplement 1.**
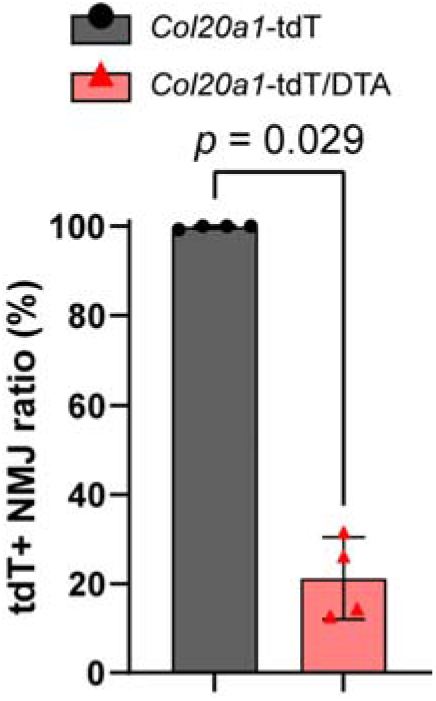
Loss of the tdTomato reporter at NMJs of *Col20a1*-tdT/DTA mice. Quantification of the tdTomato-positive (tdT+) NMJ ratio in *Col20a1*-tdT control and *Col20a1*-tdT/DTA mice at 4 weeks of age, following tamoxifen administration at P10. Nearly all NMJs retain the tdTomato reporter in control mice, whereas this proportion is significantly reduced in *Col20a1*-tdT/DTA mice. Data are presented as mean ± SD (n = 4 mice per group). Statistical significance was determined using the Mann-Whitney test. Exact *p*-values are shown for comparisons that reached statistical significance (*p* < 0.05).

